# Study on the mechanism of action of the Pt(IV) complex with lonidamine ligands by ultrafast chemical proteomics

**DOI:** 10.1101/2024.12.02.626306

**Authors:** Ekaterina A. Imaikina, Ivan I. Fedorov, Daria D. Emekeeva, Elizaveta M. Kazakova, Leila A. Garibova, Mark V. Ivanov, Ilya A. Shutkov, Alexey A. Nazarov, Mikhail V. Gorshkov, Irina A. Tarasova

**Affiliations:** V.L. Talrose Institute for Energy Problems of Chemical Physics, N.N. Semenov Federal Research Center for Chemical Physics, Russian Academy of Sciences, Leninsky Pr. 38, Bld.2, 119334, Moscow, Russia; Department of Chemistry, M.V. Lomonosov State University, Leninskie Gory 1/3, 119991, Moscow, Russia; National Research University Higher School of Economics (HSE University), Miasnitskaya Str. 20, Moscow, Russian Federation

**Keywords:** platinum complexes, dual action antitumor therapy, mechanism of action, thermal proteome profiling, expression proteomics, ultrafast proteomics

## Abstract

Platinum (II) complexes such as cisplatin among the few others are well-known and approved for clinical use as anticancer metal-based drugs. In spite of their successful and wide acceptance, the respective chemotherapy is associated with severe side effects and the ability of tumors to quickly develop resistance. To overcome these drawbacks the novel strategy is considered, which is based on the use of platinum complexes with bioactive ligands attached to act in synergy with platinum and further improve its pharmacological properties. Among the recently introduced such multi-action prodrugs is Pt(IV) complex with two lonidamine ligands, the latter selectively inhibiting hexokinase and, thus, the glycolysis in cancer cells. While platinum based multi-action prodrugs are exhibiting increased levels of activity towards cancer cells and, thus, considered as potent to overcome the resistance to cisplatin, there is a crucial need to uncover their mechanism of action by revealing all possibly affected processes and targets across the whole cellular proteomes. These are the challenging tasks in proteomics requiring high-throughput analysis of hundreds of samples for just a single drug-to-proteome system. In this work we performed these analyses for 8-azaguanine and experimental Pt(IV)-lonidamine complex applied to ovarian cancer cell line A2780, using both mechanism- and compound-centric chemical proteomics approaches based on ultrafast expression proteomics and thermal proteome profiling, respectively. Analysis of data obtained for Pt(IV)-lonidamine complex revealed regulation of proteins involved in glucose metabolic process associated with lonidamine further supporting the multi-action mechanism of this prodrug action.

## INTRODUCTION

Currently, chemotherapy is one of the most commonly used approaches for cancer treatment. Platinum-based compounds remain the leading metal-based anticancer drugs following the discovery of the anticancer activity of cisplatin by Rosenberg et al.^1–4^ Since the introduction of cisplatin, numerous platinum compounds have been studied in search for antitumor alternatives that do not exhibit the same high level of systemic toxicity and/or drug resistance.^5^ In particular, Pt(IV) complexes with biologically active ligands hold promise, as they are likely to cause fewer side effects and improve pharmacological properties of Pt(II) derivatives.^6–9^ Specifically, Pt(IV) compounds function as prodrugs, that is, they are activated by reduction primarily within cancer cells to release the original Pt(II) drug along with the two axial ligands.^10^ The octahedral coordination geometry of Pt(IV) complexes provides a variety of ligand options, which can enhance the activity and specificity of the drug, as well as exhibit their own activity.^9^ In particular, it was found that the antitumor effect can be ameliorated by incorporating a cancer cell-specific targeting moiety into Pt(IV) prodrugs,^11–13^ such as lonidamine, which specifically inhibits hexokinase in solid tumors.^14^ By incorporating two lonidamine compounds as axial ligands in the platinum(IV) complex, a combination of [*cis*-ethane-1,2-diaminedichloridoplatinum (II)] with lonidamine at both axial positions (Pt(IV)-lonidamine complex) has been introduced recently,^15^ it demonstrated increased toxicity towards a variety of cancer cell lines compared to other Pt(IV) prodrugs in in vitro studies.^16,17^ However, the comprehensive map of possible mechanisms of action has yet to be revealed, so as the drug targets at the whole proteome level.

In recent years, chemical proteomics has emerged as an instrument for proteome-wide studying of biochemical processes within cells activated by drug treatment.^18,19^ It is based on quantitative proteomic analysis of protein expression in response to the treatment using high-performance liquid chromatography coupled with tandem mass spectrometry (HPLC–MS/MS), further providing new insights into the mechanisms of chemotherapeutic agent action and the identification of therapeutic targets.^20^ Among the chemoproteomic approaches, thermal proteome profiling (TPP) is currently the method of choice for identifying the drug targets and in-depth examination of drug-target interactions on a proteome-wide scale.^21–23^ TPP allows uncovering the so-called off-targets of a drug interaction which give more complete picture of the drug action mechanisms, crucially important at the stage of novel drug development and/or repurposing.^24^ However, current TPP protocols require proteome-wide analyses of multiple temperature fractions and, even more in 2D-TPP realization, when both temperature and drug concentration change. This limits a variety of samples accessible for quantitative analysis. The problem is more acute in case of multi-action drugs, such as, for example, Pt(IV)-two ligand complexes when the possible variety of ligands creates a whole spectrum of antitumor candidates, each exhibiting unique mechanisms of action and, thus, requiring careful examination and identification both targets and off-targets across a particular cellular proteome.^9^ Needless to say, the proper quantitative proteomic methodology requires analysis of each proteome in several biological and sample replicates that further aggravates the situation.

In summary, while the concept of multi-action drugs opens up a lot of space for the development of new antitumor agents, the contribution to these developments from proteomics is limited by enormously high analysis time requirements and expenses (e.g., due to extensive use of labeling techniques in employed TMT protocols). Among the recent attempts to overcome the above limitation is the use of the so-called ultrafast proteomics approaches.^25,26^ MS/MS-free method DirectMS1^27^ is among these approaches, which was applied recently in a number of chemical proteomics studies, including TPP for identification of drug targets and associated cellular processes.^28,29^ In this work, we have combined the capabilities of both expression proteomics and TPP, based on ultrafast proteome analysis, to obtain the list of possible targets of two anti-tumor drugs, a well-known and extensively characterized 8-azaguanine used as a control for the developed workflow and Pt(IV)-lonidamine complex.

## METHODS

### Cell culture

The human A2780 ovarian carcinoma cell line was obtained from the European collection of authenticated cell cultures (ECACC; Salisbury, UK). Cells were grown in RPMI-1640 medium (Capricorn Scientific GmbH, Germany) supplemented with L-glutamine and 10 % fetal bovine serum (Gibco™, Brazil). The cells were cultured at 37 °C in a humidified 5 % CO_2_ atmosphere and subcultured 2 times a week.

### Drugs

Two anti-tumor drugs were used in the study: triazolo[4,5-d]pyrimidine 8-azaguanine and (OC-6–33)-dichloridobis[1-(2,4-dichlorobenzyl)-1H-indazole-3-carboxylato](ethane-1,2-diamine)platinum(IV) (Pt(IV)-lonidamine complex).^15^ The former one is a purine analog exerting cytotoxic activity by inhibiting purine nucleoside phosphorylase (PNPase),^30^ and the latter has been introduced recently as a dual-action prodrug described elsewhere.^17^ The chemical structures of these drugs are shown schematically in Figure 1.

**Figure 1.**
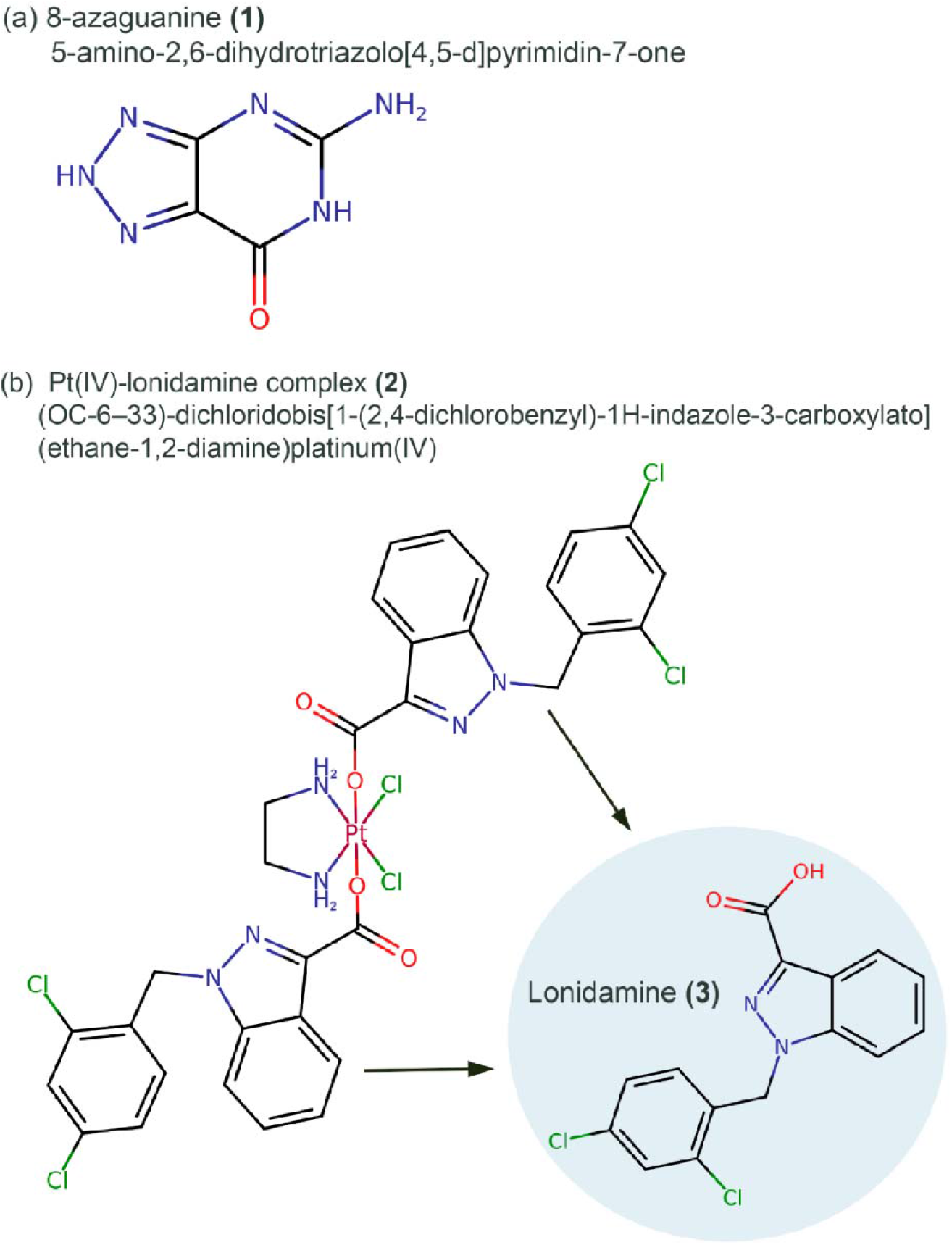
Investigated compounds with antitumor activity: (a) 8-azaguanine (**1**) which is the triazole analogue of guanine; and (b) (**2**) complex of platinum (IV) with two lonidamine (**3**) ligands.

### Sample preparation for expression proteomics

After incubation in individual culture flasks, the cells were treated with (**2**) dissolved in DMSO at 0.05% concentration, for 72 hours. For the cells’ treatment, the set of drug concentrations was 0.0078125, 0.03125, 0.0625, 0.125, 0.25, 0.5 and 1 μM, and a concentration of 0 μM (pure 0.05% DMSO treatment) was used as a negative control. Each concentration point was repeated four times. After treatment, the cells were subjected to centrifugation at 1500 rpm. The resulting cellular pellets (cells per sample) were solubilized in 100 μM lysis buffer (0.1% ProteaseMax (Promega, USA), 100 mM ABC, 10% ACN). The mixture was incubated at room temperature for one hour and then boiled for 6 minutes on a thermoshaker at 95°C, after which it was immediately cooled down on ice. To lyse the cells, the samples were placed in the ultrasonic bath for 15 minutes and underwent additional ultrasonic homogenization for 2 min (5 s on 5 s off) on 40% amplitude (QSonica Q125, Newtown, Connecticut, USA). Next, the mixture was centrifuged for 7 minutes at 10,000 g. Subsequently, a protein-containing supernatant was collected. After protein extraction, the concentration was measured by BCA kit (Thermo Scientific, Germany). For disulfide bonds reduction in protein samples, 10 mM DTT was added for 20 minutes at 56°C, followed by incubation with 20 mM Iodoacetamide (IAA) for 30 min in the dark at room temperature. For protein digestion, trypsin (Sequencing Grade Modified Trypsin, Promega, USA) was used at a ratio of 1:50 (w/w) with protein mixtures. The samples were then incubated at 37°C for 18 hours. To stop the digestion, 1% TFA was added. Samples were then centrifuged for 15 minutes at 20,000g. A supernatant was collected and the peptide samples were subsequently dried in a vacuum concentrator.

### Sample preparation for TPP

200×10^6^ A2780 cells were resuspended in a 2.5 ml lysis buffer (1 ml protease inhibitor 25x (cOmplete™ EDTA-free Protease Inhibitor Cocktail, Roche, Switzerland), 1.5 ml 1x PBS). Solution was divided into 10 aliquots and cells were lysed with 5 freeze-thaw cycles (2 min in liquid nitrogen 2 min at 25 °C). Samples were centrifuged for 20 min on 20000 *g*, then supernatant was taken, and aliquots were combined together. Protein concentration was measured by BCA kit (Thermo Scientific, Germany). The stock solution was divided into control (Control) and treatment (Treatment) samples and PBS solution was added to reach the final protein concentration of 2 ug/ul. Test sample for Pt(IV)-lonidamine complex was treated with 20 uM solution in DMSO 1:100 (v/v) and the control sample was treated with DMSO 1:100 v/v. For 8-Azaguanine, the sample was treated with 20 mM drug’s solution in DMSO at the ratio of 1:32 (v/v), and the control sample was treated with DMSO at the same ratio of 1:32 (v/v). All samples were incubated for 2 hours at 25 °C, followed by supernatant extraction and protein concentration measurements as described above. Then, both treatment and control samples were each divided into 10 groups of 5 aliquots. Each group was incubated for 3 min at one of the following temperatures: 37 °C, 41 °C, 44 °C, 47 °C, 50 °C, 53 °C, 56 °C, 59 °C, 63 °C, 67 °C. Samples were cooled on ice, centrifuged for 30 min at 50000g and supernatants were taken. DTT 10 mM was added for 20 min at 56°C, followed by incubation with IAA for 30 min in dark at room temperature. Dry pellets were resuspended sequentially with 40 uL 0.1% ProteaseMax (Promega, USA) in 100 mM ABC and 40 uL 100 mM ABC. Trypsin protease (Sequencing Grade Modified Trypsin, Promega, USA) was added in 1:75 (w/w) ratio, and samples incubated overnight at 37°C. Digestion was stopped by adding 1% TFA, then samples were centrifuged for 15 min at 20000 g, supernatants were taken and dried in a vacuum concentrator.

### Ultrafast LC-MS1 data collection

LC-MS data were collected at the IBMC Human Proteome Core Facility (Moscow, Russia, https://en.ibmc.msk.ru/core). The experiments were performed using an Orbitrap Q Exact HFX mass spectrometer (Thermo Scientific, San Jose, CA, USA) coupled to UltiMate 3000 LC system (Thermo Fisher Scientific, Germering, Germany). For separation, two different columns, a μ-Precolumn C18 PepMap100, 5 μm, 300 μm i.d, 5 mm length, 100 Å (Thermo Fisher Scientific, Waltham, USA) and a reverse-phase capillary column Peaky, Reprosil-Pur C18 AQ 1.9 mm, 75 μm i.d., 5 cm length, 120 Å (Molekta, Moscow, Russia) were used. The mobile phases were: (A) 0.1% formic acid (FA) in water; (B) 80% ACN, 0.1% FA in water. A gradient from 5% to 35% phase B in 4.8 min at 1.5 μl/min was applied. Data acquisition was performed using MS1 mode only. Full MS scans were acquired in the range from *m/z* 375 to 1,500, with a resolution of 120,000 at *m/z* 200 and Automatic Gain Control (AGC) setting of 4*10^5^, 1 microscan per cycle, and a maximum injection time of 50 milliseconds. The samples were resuspended in phase A, and 1 μg were loaded per injection.

### Expression proteomics data analysis

The conversion of the raw experimental data to. mzML format was performed using ThermoRawParser, v. 1.4.3.^31^ Protein identification was conducted using ms1searchpy, v.2.3.20, search engine.^27^ The SwissProt canonical human protein database (version from 25.01.2023 containing 20,389 entries) was employed for the search, augmented with obviously incorrect decoys generated by randomly rearranging amino acid sequences from the database. The following search parameters were used: enzyme - trypsin; a minimum of 3 scans per peptide ion cluster formed by the ^13^C isotope distribution of peptides; at least 2 peaks in the cluster, including the monoisotopic peak; initial mass accuracy of 8 ppm; peptides without missed cleavage sites and with lengths between 7 and 30 amino acid residues (484,765 peptides) were considered; charge states ranged from 1+ to 6+; and carboxymethylation of cysteine as fixed modification. Mass calibration was achieved by retention time analysis using DeepLC.^38^ The resulting list of protein identifications was filtered to 5% FDR using “target-decoy” approach^39^, where the decoy database was generated through a modified pseudo-shuffling of amino acid sequences from target proteins.

To identify differentially expressed proteins, a quantitative analysis was conducted using DirectMS1Quant^34^ and Diffracto^35^ softwares. Both algorithms provide an estimate of the fold change (FC) in protein abundance, as well as statistical significance of results based on a p-value adjusted for multiple comparisons using the Benjamini–Hochberg procedure.

The Diffracto algorithm has a higher sensitivity to changes in proteomes, however, this may come at the cost of a higher rate of false positives.^40^ Based on the results of the Diffacto quantification, a semi-dynamic selection of proteins that are differentially regulated was performed using QRePS software.^36^ A fixed p-value threshold was set at *FC=*1.2, and the threshold for concentration changes was calculated by considering the statistical distribution of the experimental data: *log*_2_*(FC) =<log*_2_*(FC)>±1.5σ(log*_2_*(FC))*.

Functional enrichment analysis was performed via API-STRING^37^ for a set of proteins that showed statistically significant changes in their concentration after treatment with the drug.

### TPP data analysis

The TPP data processing workflow used in this study was described elsewhere.^29^ Briefly, it has three main stages: (1) protein identification within the analyzed fractions using ms1searchpy, (2) quantitative analysis of these protein identifications with DirectMS1Quant, and (3) identification of proteins with statistically significant changes in denaturation temperature following drug treatment. Protein identification was based on the SwissProt canonical human sequences library, version dated 25.01.2023, restricted to *Homo sapiens* and containing 20,389 proteins. Thermo raw files were converted to the mzML format using ThermoRawFileParser. Identification was conducted using ms1searchpy v. 2.3.20 as described above in the “Expression proteomics data analysis” section.

#### Solubility curve construction

The result of protein denaturation is the subsequent decrease in its relative concentrations in the higher temperature fractions. For normalization of the quantitation data, we, first, assume that each identified protein has a maximum concentration in a 37°C fraction. Generally speaking, this may not always hold true in some limited number of cases. Then, for each of the 10 temperature points in the quantitation data, a median relative concentration based on the intensities of protein’s peptides as described elsewhere^34^ is calculated from all 5 technical replicates. Then, for each temperature point, the fold changes of protein concentrations compared with the one in the 37 °C fraction were estimated by DirectMS1Quant software, and only proteins with a statistically significant FC at least at one temperature point were considered. In other words, if the concentration of a protein did not change in all fractions, this protein was disqualified from further analysis. Then, the experimental data obtained for remaining proteins were approximated by the theoretical solubility curves defined as follows:

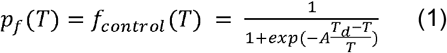

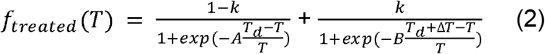

in which *T*_*d*_ is the protein melting temperature, *A* and *B* are the protein’s thermodynamic coefficients and *k* is the efficiency of drug-to-protein binding reaction as described earlier.^29,41^ Δ*T*_*d*_ is the change in melting temperature after treatment and the measured value used further for identifying drug target candidates. All proteins with the accuracy of R^2^<0.8 for the experimental data fit by theoretical solubility curves were further disqualified from the analysis. At the final step of the analysis, the distribution of qualified proteins by Δ*T*_*d*_ was determined. This distribution is normal as expected for the majority of qualified proteins, and the ones with the change in melting temperature exceeding 3σ were considered as the outliers with 99.7% of the probability (three-sigma rule of thumb) and considered as drug target candidates. Among them, we selected only the ones with reaction coefficient efficiency exceeding 50% for the short list of drug targets. Note that this assumption brings some ambiguity in the results as even weak interactions between a drug and a particular protein may affect the protein functioning, thus being too conservative. Pipelines for both expression proteomics and the TPP method are briefly presented in Figure 2, while a more detailed pipeline for TPP is shown in Figure S1 and mass spectrometry proteomics experimental data has been deposited to the ProteomeXchange Consortium via the PRIDE partner repository with the dataset identifier PXD058413.

**Figure 2.**
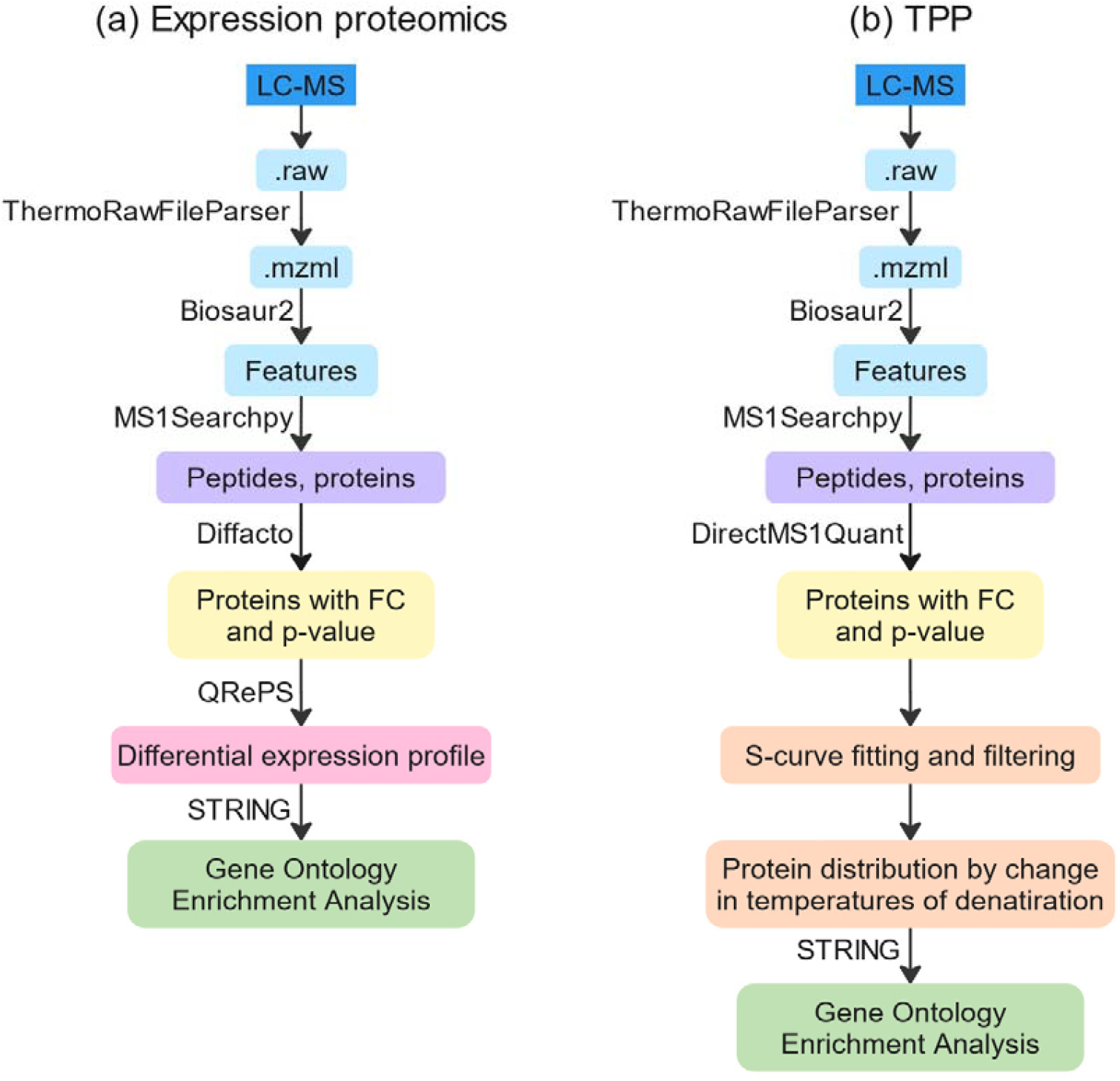
Workflow for data processing used in this work: (a) expression proteomics; (b) TPP approach. In both cases the raw LC-MS data was converted into .mzml format using a ThermoRawFileParser.^31^ Biosaur^32^ was employed for feature detection in MS1 spectra, followed by protein identification using ms1searchpy engine.^33^ Quantitative analysis was performed on the expression proteomics samples using the DirectMS1Quant^34^ and Diffacto^35^ algorithms. A differential expression profile was then selected using QRePS^36^, based on which gene ontologies were analyzed using the STRING database.^37^ With regard to TTP probes, S-shape curve approximations for the protein solubility curves were generated to identify drug targets based on the measured changes in protein melting temperatures after the cell treatment. The combined (TPP+expression proteomics) interactome maps were built using StringDB, v.12 software, as well.

## RESULTS AND DISCUSSION

First, we started with analysis of the results obtained by expression proteomics. Quantitative comparison of control vs. treatment proteomes was performed using Diffacto software to identify a group of proteins with a statistically significant change in expression after the treatment. Over 250 and 100 differentially expressed proteins (DEPs) in treated samples were identified for the 8-azaguanine and Pt(IV)-lonidamine complex concentrations of 3.125 μM and 0.0625 μM, respectively. These concentrations are close to these drugs’ corresponding IC_50_ values of 3.6 ± 0.6 μM and 0.07±0.02 μM estimated using the MTT assay for the metabolic activity of A2780 cells under study as shown in Figure S2 in the Supplemental Materials.

Generally, the experiment was conducted by treating cells with eight different concentrations of each drug. Using the GO analysis results, the biological processes that were most enriched in cells in response to the drug treatment are sketched at Figure S3. Under the influence of 8-azaguanine, in a wide variety of concentrations, the proteins involved in glycolysis and nucleotide metabolism, particularly purine metabolism, significantly alter their concentration. This finding is consistent with 8-azaguanine’s known mechanism of action as an antimetabolite that resembles the purine base guanine^42^, and reportedly the polyploid-specific disruptor of proliferation as suggested recently.^43^ In the case of the Pt(IV)-lonidamine complex, there are observed changes in the processes of glycolysis. These processes may be influenced by lonidamine, which is a component of the complex that inhibits glycolysis.^44^ Furthermore, we observe enrichment in some DNA-related processes. This effect is expected due to the platinum core of the complex.^45^

Second, we analyzed the results of TPP data. Between 1000 and 1200 proteins in each temperature fraction were identified in both ligands. 434 proteins for 8-azaguanine and 288 proteins for Pt(IV)-lonidamine complex were identified in all temperature fractions that passed the above criteria for the accuracy of S-shaped curve fitting of experimental data. The standard deviation for distribution of ΔT was found, and the proteins that passed the selection criteria for the statistical significance of the melting temperature change were those whose temperature was shifted in one, or the other direction by more than 3σ.

Analysis of the distributions of qualified proteins by changes in melting temperatures has resulted in 22 and 15 proteins for 8-azaguanine and Pt(IV)-lonidamine complex with melting temperature shifts exceeding the. Solubility curves for all target proteins are provided in Figure S4 of Supplemental Information. Enrichment analysis by cellular functions and components for proteins identified as topotecan targets was performed using gene ontologies (GO) as well. The results of the analysis of the 8-azaguanine reveals that the some identified proteins take part in processes such as: RNA-related - HNRC1, HNRC3, HNRC4, RM04 (mitochondrial), CLU (mitochondrial), RT27, TRM6 (tRNA, mRNA), EIF3A (mRNA, eukaryotic translation initiation factor 3 complex), CLH1 (ds-RNA, RNA), EFGM (translation elongation, mitochondrial), LPPRC (nuclear mRNA export), RPAC1 (DNA-dependent RNA polymerases subunit); Ubiquitin-related - PLAP, PDC6I (ubiquitin-independent protein catabolic process via the multivesicular body sorting pathway), OTU1, CAN1 (catalyzes limited proteolysis of substrates involved in cytoskeletal remodeling); Transport - GOPC (Plays a role in intracellular protein trafficking and degradation). This observation is in good agreement with the known mechanism of 8-azaguanine. And results analysis for the Pt(IV)-lonidamine complex reveal that the identified proteins take part in processes such as: Glucose 6-phosphate metabolic process - HXK1, HXK2, PFKAM, ODPB (glucose metabolic process); RNA-related - SYEP, ZFP62, SYFB; Zinc/iron/metal ion binding: P3H1, PLOD1, SYEP, ZFP62. Also, hexokinase-2 and hexokinase-1, two of the targets, are direct targets of lonidamine^46^; accordingly, glucose 6-phosphate metabolic process is an expected process in the mechanism of action of this drug.

For the interactomes DEPs identified in expression proteomics (for concentration of 3.125 μM and 0.0625 μM, respectively) analysis were combined with the targets from TPP study. This combination allows revealing the networks indirectly involved in the cell’s response to the treatment due to the drug binding to a protein not showing as being differentially expressed, yet changing its functionality in the cell. For the combined set an interaction map was generated using StringDB with the confidence level setting of 0.7 (Figure 3).

**Figure 3.**
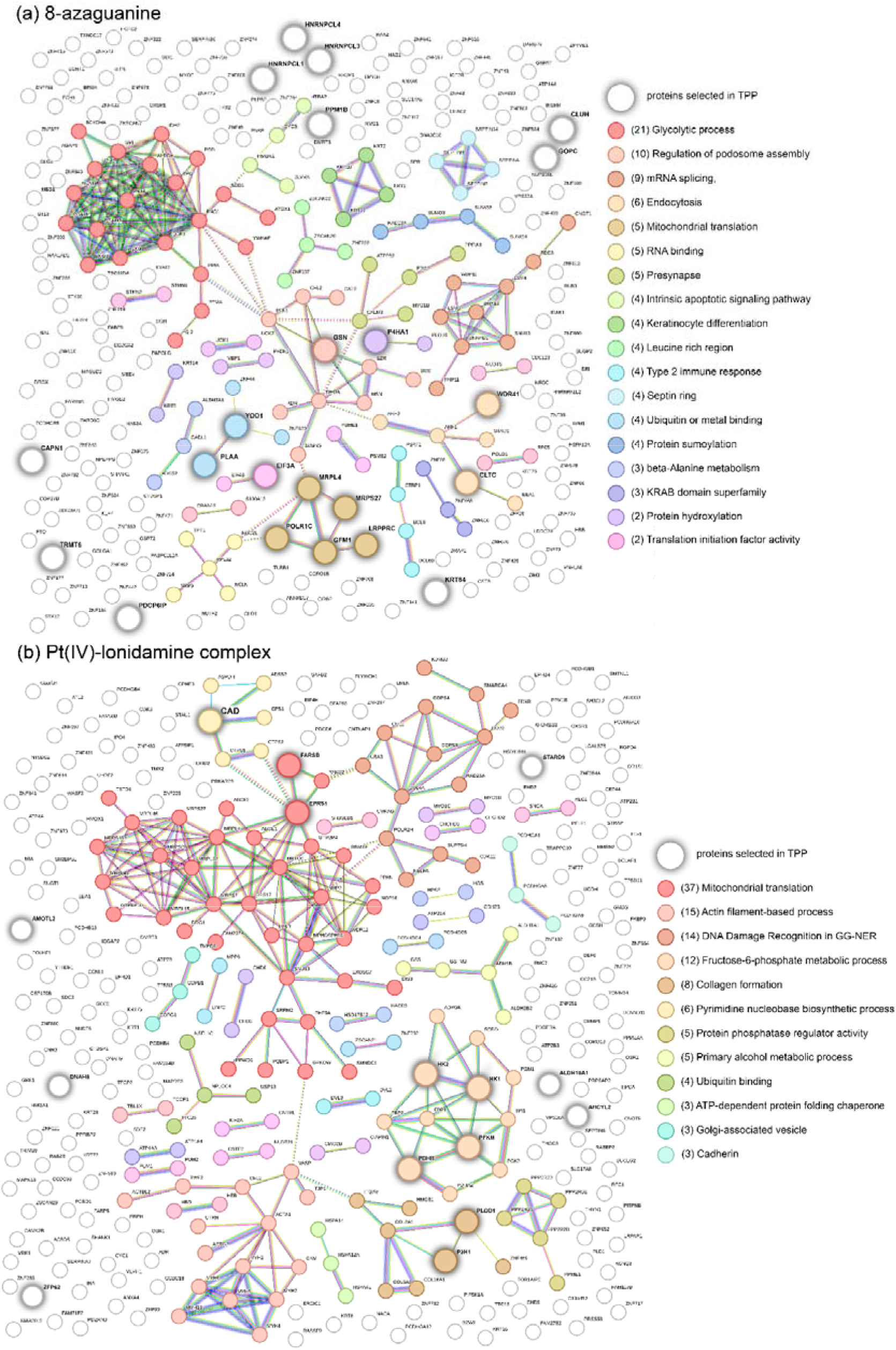
Cluster analysis of interactions between differentially regulated proteins from expression proteomic data and direct drug targets identified in TPP for 8-azaguanine (a) and prodrug (b). StringDB was used for the interactomes’ mapping with the minimum required interaction score of 0.7 (high confidence), the number of clusters for the analysis was 26 and 31 respectively. The interacting clusters are highlighted in different colors. Proteins identified by TPP as targets are shown in larger sizes.

Among the proteins that showed changes in their concentration after treatment with 8-azaguanine, distinct groups of proteins involved in glycolysis, mRNA splicing, mitochondrial translation, and RNA binding. 8-Azaguanine, as discussed above, is expected to affect these processes due to its mechanism of action as an antimetabolite that interferes with nucleotide synthesis.

The interactome map for the Pt(IV)-lonidamine complex shown in Fig.3b includes a cluster of glycolytic proteins with hexokinases identified as targets by TPP. Importantly, hexokinase is reported in the literature as a potential target for lonidamine. Among the processes listed in Table 2, the metabolism of fructose 6-phosphate is highlighted. This process is directly linked to the first glycolysis reaction, which is catalyzed by the hexokinase exactly. In addition, there is apparently a large cluster of proteins involved in mitochondrial translation, which were found regulated from expression proteomics. This is also consistent with the findings that lonidamine strongly affects mitochondrial function.^44^ The cluster of proteins involved in the process of DNA damage recognition is of particular interest in the map on Fig.3b. Platinum-based compound, which is the part of experimental prodrug used in this study is causing irreversible damage to DNA through the mechanism of action similar to the one of cisplatin. Changes in the expression of DNA repair proteins indicate that the Pt(IV)-lonidamine prodrug also alters the structure of DNA, thus further supporting the expected multi-action mechanism from the synergetic effect of both lonidamine ligands and platinum complexes. Target proteins and the ones regulated by treatment identified from TPP experiments and expression proteomics analysis, respectively, are listed for both drugs in Supplemental Table S1.

## CONCLUSIONS

The ultrafast method of quantitative MS1-only proteome-wide analysis was used to study interaction of chemotherapy drugs with cancer cell lines using two chemical proteomics approaches. The first one was a compound-centered approach based on thermal proteome profiling, and the other one was a mechanism-centered approach based on expression proteomics. These approaches were implemented for identifying mechanisms of action and targets of an experimental prodrug currently under development, the Pt(IV)-contained complex with two lonidamine ligands, applied to ovarian cancer cell line A2780. The workflow used in this work was combining both TPP and expression proteomics results to reveal intracellular processes regulated by the treatment. The anticancer drug 8-azaguanine with the known main mechanisms of action, which include the interference with nucleotide synthesis and antimetabolic activity was used to validate the workflow. The results obtained for 8-azaguanine confirmed its main mechanism of action revealing ribonucleoside, ADP, and ATP metabolic processes regulated by this drug treatment. For the experimental prodrug the study revealed regulation of processes known as associated with both platinum-contained drugs and lonidamine. Specifically, among the top enriched processes were the ones related to DNA damage and repair associated with platinum and lonidamine-induced glucose metabolism involving hexokinase, the latter identified by TPP as the prodrug’s substrate.The results obtained for the experimental Pt(IV)-contained complex with lonidamine support its dual-action mechanism and potential as a new generation antitumor drug with increased cytotoxicity towards cancer cells.

## ABBREVIATIONS

ADP: Adenosine Diphosphate
ATP: Adenosine Triphosphate
CAN1: Calcium and Integrin Binding Protein 1
CLH1: Chloride Channel Protein 1
CLU: Clusterin
DNA: Deoxyribonucleic Acid
EFGM: Elongation Factor G Mitochondrial
EIF3A: Eukaryotic Translation Initiation Factor 3 Subunit A
FC: Fold Change
FDR: False discovery rate
GOPC: Golgi Associated PDZ and Coiled Coil Motif Containing Protein
HNRC1: Heterogeneous Nuclear Ribonucleoprotein C1
HNRC3: Heterogeneous Nuclear Ribonucleoprotein C3
HNRC4: Heterogeneous Nuclear Ribonucleoprotein C4 HXK1 Hexokinase 1
HXK2: Hexokinase 2
LPPRC: Leucine-Rich Pentatricopeptide Repeat Containing Protein
mRNA: Messenger Ribonucleic Acid
MS1: Mass Spectrometry, level 1
ODPB: Pyruvate Dehydrogenase E1 Component Subunit Beta
OTU1: Ovarian Tumor Domain-Containing Protein 1
P3H1: Prolyl 3 Hydroxylase 1
PDC6I: Programmed Cell Death 6 Interacting Protein
PFKAM: Phosphofructokinase Muscle Type
PLAP: Placental Alkaline Phosphatase
PLOD1: Procollagen Lysine Hydroxylase 1
Pt(IV): Platinum in oxidation state IV
TPP: Thermal Proteome Profiling
DMSO: Dimethyl Sulfoxide
ABC: Ammonium Bicarbonate
MS/MS: Tandem Mass Spectrometry
HPLC: High-Performance Liquid Chromatography
FA: Formic Acid
ACN: Acetonitrile
RPMI: Roswell Park Memorial Institute Medium
IC_50_: Half-maximal inhibitory concentration
DEPs: Differentially Expressed Proteins
GO: Gene Ontology
PBS: Phosphate Buffered Saline
BCA: Bicinchoninic Acid Assay
CO_2_: Carbon Dioxide
MTT: Tetrazolium Reduction Test
RM04: Mitochondrial Ribosomal Protein L4
RNA: Ribonucleic Acid
RPAC1: RNA Polymerase I Subunit A
RT27: Retinitis Pigmentosa GTPase Regulator-Interacting Protein 1 (RPGRIP1)
SYEP: Seryl tRNA Synthetase Cytoplasmic
SYFB: Phenylalanyl tRNA Synthetase Beta Subunit
TMT: Tandem mass tag
TRM6: tRNA Methyltransferase 6
ZFP62: Zinc Finger Protein 62

## AUTHORS’ CONTRIBUTION

I.I.F., D.D.E., E.M.K., and L.A.G. performed proteomic sample preparations and experiments, M.V.G. and I.A.T. were involved in planning and supervised the work on proteomics at all stages, I.A.S. and A.A.N. performed cell work, drug design, MTT tests, and cell treatment, E.A.I. and I.I.F. performed proteomic data analysis, drafted the manuscript and designed the figures, M.V.I, I.I.F, and E.M.K. developed software development and testing.for quantitative analysis, E.A.I. I.I.F., M.V.G., A.A.N., and I.A.T. aided in interpreting the results, E.A.I., I.I.F., and M.V.G. wrote and reviewed the manuscript. All authors discussed the results and commented on the manuscript.

## REFERENCES

(1) Rosenberg, B.; Vancamp, L.; Trosko, J. E.; Mansour, V. H. Platinum Compounds: A New Class of Potent Antitumour Agents. Nature 1969, 222 (5191), 385–386. 10.1038/222385a0.

(2) Brabec, V.; Hrabina, O.; Kasparkova, J. Cytotoxic Platinum Coordination Compounds. DNA Binding Agents. Coord. Chem. Rev. 2017, 351, 2–31. 10.1016/j.ccr.2017.04.013.

(3) Wheate, N. J.; Walker, S.; Craig, G. E.; Oun, R. The Status of Platinum Anticancer Drugs in the Clinic and in Clinical Trials. Dalton Trans. 2010, 39 (35), 8113–8127. 10.1039/C0DT00292E.

(4) Florea, A.-M.; Büsselberg, D. Cisplatin as an Anti-Tumor Drug: Cellular Mechanisms of Activity, Drug Resistance and Induced Side Effects. Cancers 2011, 3 (1), 1351– 1371. 10.3390/cancers3011351.

(5) Wang, D.; Lippard, S. J. Cellular Processing of Platinum Anticancer Drugs. Nat. Rev. Drug Discov. 2005, 4 (4), 307–320. 10.1038/nrd1691.

(6) Wang, Z.; Deng, Z.; Zhu, G. Emerging Platinum(IV) Prodrugs to Combat Cisplatin Resistance: From Isolated Cancer Cells to Tumor Microenvironment. Dalton Trans. 2019, 48 (8), 2536–2544. 10.1039/C8DT03923B.

(7) Okulova, Y. N.; Zenin, I. V.; Shutkov, I. A.; Kirsanov, K. I.; Kovaleva, O. N.; Lesovaya, E. A.; Fetisov, T. I.; Milaeva, E. R.; Nazarov, A. A. Antiproliferative Activity of Pt(IV) Complexes with Lonidamine and Bexarotene Ligands Attached via Succinate-Ethylenediamine Linker. Inorganica Chim. Acta 2019, 495, 119010. 10.1016/j.ica.2019.119010.

(8) Kenny, R. G.; Marmion, C. J. Toward Multi-Targeted Platinum and Ruthenium Drugs—A New Paradigm in Cancer Drug Treatment Regimens? Chem. Rev. 2019, 119 (2), 1058–1137. 10.1021/acs.chemrev.8b00271.

(9) Gibson, D. Multi-Action Pt(IV) Anticancer Agents; Do We Understand How They Work? J. Inorg. Biochem. 2019, 191, 77–84. 10.1016/j.jinorgbio.2018.11.008.

(10) Gibson, D. Platinum(IV) Anticancer Prodrugs – Hypotheses and Facts. Dalton Trans. 2016, 45 (33), 12983–12991. 10.1039/C6DT01414C.

(11) Ang, W. H.; Khalaila, I.; Allardyce, C. S.; Juillerat-Jeanneret, L.; Dyson, P. J. Rational Design of Platinum(IV) Compounds to Overcome Glutathione-S-Transferase Mediated Drug Resistance. J. Am. Chem. Soc. 2005, 127 (5), 1382–1383. 10.1021/ja0432618.

(12) Karges, J.; Yempala, T.; Tharaud, M.; Gibson, D.; Gasser, G. A Multi-Action and Multi-Target RuII–PtIV Conjugate Combining Cancer-Activated Chemotherapy and Photodynamic Therapy to Overcome Drug Resistant Cancers. Angew. Chem. Int. Ed. 2020, 59 (18), 7069–7075. 10.1002/anie.201916400.

(13) Petruzzella, E.; Sirota, R.; Solazzo, I.; Gandin, V.; Gibson, D. Triple Action Pt(IV) Derivatives of Cisplatin: A New Class of Potent Anticancer Agents That Overcome Resistance. Chem. Sci. 2018, 9 (18), 4299–4307. 10.1039/C8SC00428E.

(14) Nath, K.; Guo, L.; Nancolas, B.; Nelson, D. S.; Shestov, A. A.; Lee, S.-C.; Roman, J.; Zhou, R.; Leeper, D. B.; Halestrap, A. P.; Blair, I. A.; Glickson, J. D. Mechanism of Antineoplastic Activity of Lonidamine. Biochim. Biophys. Acta 2016, 1866 (2), 151– 162. 10.1016/j.bbcan.2016.08.001.

(15) Nosova, Y. N.; Foteeva, L. S.; Zenin, I. V.; Fetisov, T. I.; Kirsanov, K. I.; Yakubovskaya, M. G.; Antonenko, T. A.; Tafeenko, V. A.; Aslanov, L. A.; Lobas, A. A.; Gorshkov, M. V.; Galanski, M. S.; Keppler, B. K.; Timerbaev, A. R.; Milaeva, E. R.; Nazarov, A. A. Enhancing the Cytotoxic Activity of Anticancer Pt-IV Complexes by Introduction of Lonidamine as an Axial Ligand. Eur. J. Inorg. Chem. 2017, 2017 (12), 1785–1791. 10.1002/ejic.201600857.

(16) Nosova, Y. N.; Zenin, I. V.; Maximova, V. P.; Zhidkova, E. M.; Kirsanov, K. I.; Lesovaya, E. A.; Lobas, A. A.; Gorshkov, M. V.; Kovaleva, O. N.; Milaeva, E. R.; Galanski, M.; Keppler, B. K.; Nazarov, A. A. Influence of the Number of Axial Bexarotene Ligands on the Cytotoxicity of Pt(IV) Analogs of Oxaliplatin. Bioinorg. Chem. Appl. 2017, 2017 (1), 4736321. 10.1155/2017/4736321.

(17) Kasparkova, J.; Kostrhunova, H.; Novohradsky, V.; Ma, L.; Zhu, G.; Milaeva, E. R.; Shtill, A. A.; Vinck, R.; Gasser, G.; Brabec, V.; Nazarov, A. A. Is Antitumor Pt(IV) Complex Containing Two Axial Lonidamine Ligands a True Dual-or Multi-Action Prodrug? Met. Integr. Biometal Sci. 2022, 14 (7), mfac048. 10.1093/mtomcs/mfac048.

(18) Fedorov, I. I.; Lineva, V. I.; Tarasova, I. A.; Gorshkov, M. V. Mass Spectrometry-Based Chemical Proteomics for Drug Target Discoveries. Biochem. Biokhimiia 2022, 87 (9), 983–994. 10.1134/S0006297922090103.

(19) Meissner, F.; Geddes-McAlister, J.; Mann, M.; Bantscheff, M. The Emerging Role of Mass Spectrometry-Based Proteomics in Drug Discovery. Nat. Rev. Drug Discov. 2022, 21 (9), 637–654. 10.1038/s41573-022-00409-3.

(20) Saei, A. A.; Sabatier, P.; Tokat, Ü.G.; Chernobrovkin, A.; Pirmoradian, M.; Zubarev, R. A. Comparative Proteomics of Dying and Surviving Cancer Cells Improves the Identification of Drug Targets and Sheds Light on Cell Life/Death Decisions. Mol. Cell. Proteomics MCP 2018, 17 (6), 1144–1155. 10.1074/mcp.RA118.000610.

(21) Mateus, A.; Kurzawa, N.; Perrin, J.; Bergamini, G.; Savitski, M. M. Drug Target Identification in Tissues by Thermal Proteome Profiling. Annu. Rev. Pharmacol. Toxicol. 2022, 62, 465–482. 10.1146/annurev-pharmtox-052120-013205.

(22) Le Sueur, C.; Hammarén, H. M.; Sridharan, S.; Savitski, M. M. Thermal Proteome Profiling: Insights into Protein Modifications, Associations, and Functions. Curr. Opin. Chem. Biol. 2022, 71, 102225. 10.1016/j.cbpa.2022.102225.

(23) Sauer, P.; Bantscheff, M. Thermal Proteome Profiling for Drug Target Identification and Probing of Protein States. Methods Mol. Biol. Clifton NJ 2023, 2718, 73–98. 10.1007/978-1-0716-3457-8_5.

(24) Palve, V.; Liao, Y.; Remsing Rix, L. L.; Rix, U. Turning Liabilities into Opportunities: Off-Target Based Drug Repurposing in Cancer. Semin. Cancer Biol. 2021, 68, 209– 229. 10.1016/j.semcancer.2020.02.003.

(25) Messner, C. B.; Demichev, V.; Wang, Z.; Hartl, J.; Kustatscher, G.; Mülleder, M.; Ralser, M. Mass Spectrometry-Based High-Throughput Proteomics and Its Role in Biomedical Studies and Systems Biology. Proteomics 2023, 23 (7–8), e2200013. 10.1002/pmic.202200013.

(26) Fedorov, I. I.; Protasov, S. A.; Tarasova, I. A.; Gorshkov, M. V. Ultrafast Proteomics. Biochem. Mosc. 2024, 89 (8), 1349–1361. 10.1134/S0006297924080017.

(27) Ivanov, M. V.; Bubis, J. A.; Gorshkov, V.; Abdrakhimov, D. A.; Kjeldsen, F.; Gorshkov, M. V. Boosting MS1-Only Proteomics with Machine Learning Allows 2000 Protein Identifications in Single-Shot Human Proteome Analysis Using 5 Min HPLC Gradient. J. Proteome Res. 2021, 20 (4), 1864–1873. 10.1021/acs.jproteome.0c00863.

(28) Solovyeva, E. M.; Bubis, J. A.; Tarasova, I. A.; Lobas, A. A.; Ivanov, M. V.; Nazarov, A.; Shutkov, I. A.; Gorshkov, M. V. On the Feasibility of Using an Ultra-Fast DirectMS1 Method of Proteome-Wide Analysis for Searching Drug Targets in Chemical Proteomics. Biochem. Biokhimiia 2022, 87 (11), 1342–1353. 10.1134/S000629792211013X.

(29) Fedorov, I. I.; Bubis, J. A.; Kazakova, E. M.; Lobas, A. A.; Ivanov, M. V.; Emekeeva, D. D.; Tarasova, I. A.; Nazarov, A. A.; Gorshkov, M. V. On the Utility of Ultrafast MS1-Only Proteomics in Drug Target Discovery Studies Based on Thermal Proteome Profiling Method. Anal. Bioanal. Chem. 2024, 416 (18), 4083–4089. 10.1007/s00216-024-05330-9.

(30) Beauchamp, L. M.; Tuttle, J. V.; Rodriguez, M. E.; Sznaidman, M. L. Guanine, Pyrazolo[3,4-d]Pyrimidine, and Triazolo[4,5-d]Pyrimidine (8-Azaguanine) Phosphonate Acyclic Derivatives as Inhibitors of Purine Nucleoside Phosphorylase. J. Med. Chem. 1996, 39 (4), 949–956. 10.1021/jm950736k.

(31) Hulstaert, N.; Shofstahl, J.; Sachsenberg, T.; Walzer, M.; Barsnes, H.; Martens, L.; Perez-Riverol, Y. ThermoRawFileParser: Modular, Scalable, and Cross-Platform RAW File Conversion. J. Proteome Res. 2020, 19 (1), 537–542. 10.1021/acs.jproteome.9b00328.

(32) Abdrakhimov, D. A.; Bubis, J. A.; Gorshkov, V.; Kjeldsen, F.; Gorshkov, M. V.; Ivanov, M. V. Biosaur: An Open-Source Python Software for Liquid Chromatography–Mass Spectrometry Peptide Feature Detection with Ion Mobility Support. Rapid Commun. Mass Spectrom. n/a (n/a), e9045. 10.1002/rcm.9045.

(33) Ivanov, M. V.; Tarasova, I. A.; Levitsky, L. I.; Solovyeva, E. M.; Pridatchenko, M. L.; Lobas, A. A.; Bubis, J. A.; Gorshkov, M. V. MS/MS-Free Protein Identification in Complex Mixtures Using Multiple Enzymes with Complementary Specificity. J. Proteome Res. 2017, 16 (11), 3989–3999. 10.1021/acs.jproteome.7b00365.

(34) Ivanov, M. V.; Bubis, J. A.; Gorshkov, V.; Tarasova, I. A.; Levitsky, L. I.; Solovyeva, E. M.; Lipatova, A. V.; Kjeldsen, F.; Gorshkov, M. V. DirectMS1Quant: Ultrafast Quantitative Proteomics with MS/MS-Free Mass Spectrometry. Anal. Chem. 2022, 94 (38), 13068–13075. 10.1021/acs.analchem.2c02255.

(35) Zhang, B.; Pirmoradian, M.; Zubarev, R.; Käll, L. Covariation of Peptide Abundances Accurately Reflects Protein Concentration Differences*. Mol. Cell. Proteomics 2017, 16 (5), 936–948. 10.1074/mcp.O117.067728.

(36) Kazakova, E. M.; Solovyeva, E. M.; Levitsky, L. I.; Bubis, J. A.; Emekeeva, D. D.; Antonets, A. A.; Nazarov, A. A.; Gorshkov, M. V.; Tarasova, I. A. Proteomics-Based Scoring of Cellular Response to Stimuli for Improved Characterization of Signaling Pathway Activity. Proteomics 2023, 23 (5), e2200275. 10.1002/pmic.202200275.

(37) Szklarczyk, D.; Kirsch, R.; Koutrouli, M.; Nastou, K.; Mehryary, F.; Hachilif, R.; Gable, A. L.; Fang, T.; Doncheva, N. T.; Pyysalo, S.; Bork, P.; Jensen, L. J.; von Mering, C. The STRING Database in 2023: Protein-Protein Association Networks and Functional Enrichment Analyses for Any Sequenced Genome of Interest. Nucleic Acids Res. 2023, 51 (D1), D638–D646. 10.1093/nar/gkac1000.

(38) Bouwmeester, R.; Gabriels, R.; Hulstaert, N.; Martens, L.; Degroeve, S. DeepLC Can Predict Retention Times for Peptides That Carry As-yet Unseen Modifications. Nat. Methods 2021, 18 (11), 1363–1369. 10.1038/s41592-021-01301-5.

(39) Elias, J. E.; Gygi, S. P. Target-Decoy Search Strategy for Increased Confidence in Large-Scale Protein Identifications by Mass Spectrometry. Nat. Methods 2007, 4 (3), 207–214. 10.1038/nmeth1019.

(40) Gabdrakhmanov, I. T.; Gorshkov, M. V.; Tarasova, I. A. Proteomics of Cellular Response to Stress: Taking Control of False Positive Results. Biochem. Mosc. 2021, 86 (3), 338–349. 10.1134/S0006297921030093.

(41) Fedorov, I. I.; Ivanov, M. V.; Gorshkov, M. V. On the Effect of Drug-to-Protein Reaction Kinetics on the Results of Thermal Proteome Profiling. Anal. Chem., submitted 2024.

(42) Nelson, J. A.; Carpenter, J. W.; Rose, L. M.; Adamson, D. J. Mechanisms of Action of 6-Thioguanine, 6-Mercaptopurine, and 8-Azaguanine. Cancer Res. 1975, 35 (10), 2872–2878.

(43) Choudhary, A.; Zachek, B.; Lera, R. F.; Zasadil, L. M.; Lasek, A.; Denu, R. A.; Kim, H.; Kanugh, C.; Laffin, J. J.; Harter, J. M.; Wisinski, K. B.; Saha, S.; Weaver, B. A.; Burkard, M. E. Identification of Selective Lead Compounds for Treatment of High-Ploidy Breast Cancer. Mol. Cancer Ther. 2016, 15 (1), 48–59. 10.1158/1535-7163.MCT-15-0527.

(44) Huang, Y.; Sun, G.; Sun, X.; Li, F.; Zhao, L.; Zhong, R.; Peng, Y. The Potential of Lonidamine in Combination with Chemotherapy and Physical Therapy in Cancer Treatment. Cancers 2020, 12 (11), 3332. 10.3390/cancers12113332.

(45) Zhang, C.; Xu, C.; Gao, X.; Yao, Q. Platinum-Based Drugs for Cancer Therapy and Anti-Tumor Strategies. Theranostics 2022, 12 (5), 2115–2132. 10.7150/thno.69424.

(46) Pelicano, H.; Martin, D. S.; Xu, R.-H.; Huang, P. Glycolysis Inhibition for Anticancer Treatment. Oncogene 2006, 25 (34), 4633–4646. 10.1038/sj.onc.1209597.

