## Supplementary material for "Study on the mechanism of action of the Pt(IV) complex with lonidamine ligands by ultrafast chemical proteomics": Figure S1,Figure S2,Figure S3,Figure S4

### **SUPPLEMENTARY MATERIALS**

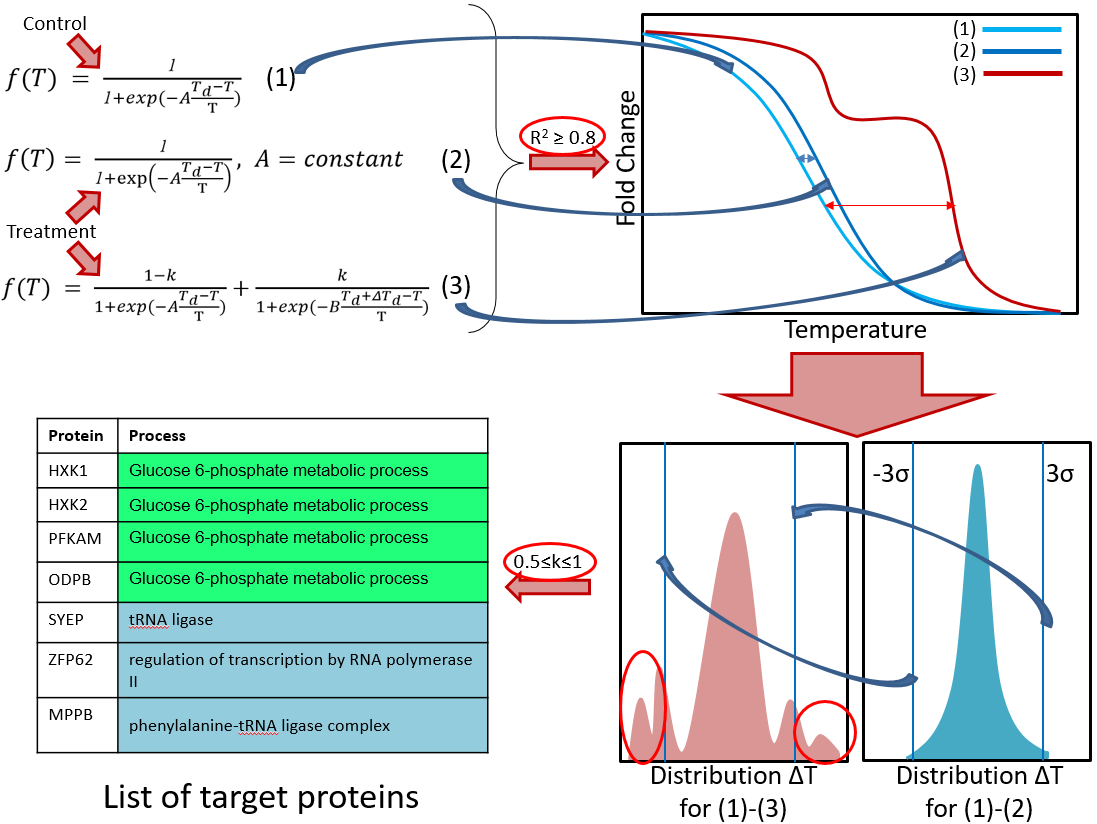

**Figure S1.** Workflow of TPP data analysis. Proteins are qualified for the final stage of the analysis and selection of targets based on the accuracy of their experimental data points fitted by S-shape solubility curves defined by Eq.1 and 2. The qualification criterion was R^2^>0.8 for the fit. Then, the distribution of qualified proteins without statistical change in ∆T by melting temperature shift was plotted to determine the standard deviation error, σ, on the assumption that most of the proteins are not experiencing the change in melting temperature, thus exhibiting normal distribution. Proteins with the melting temperature in second distribution changes exceeding 3σ are considered as the most probable drug targets. Then, targets were selected by k>0.5 to the final list.

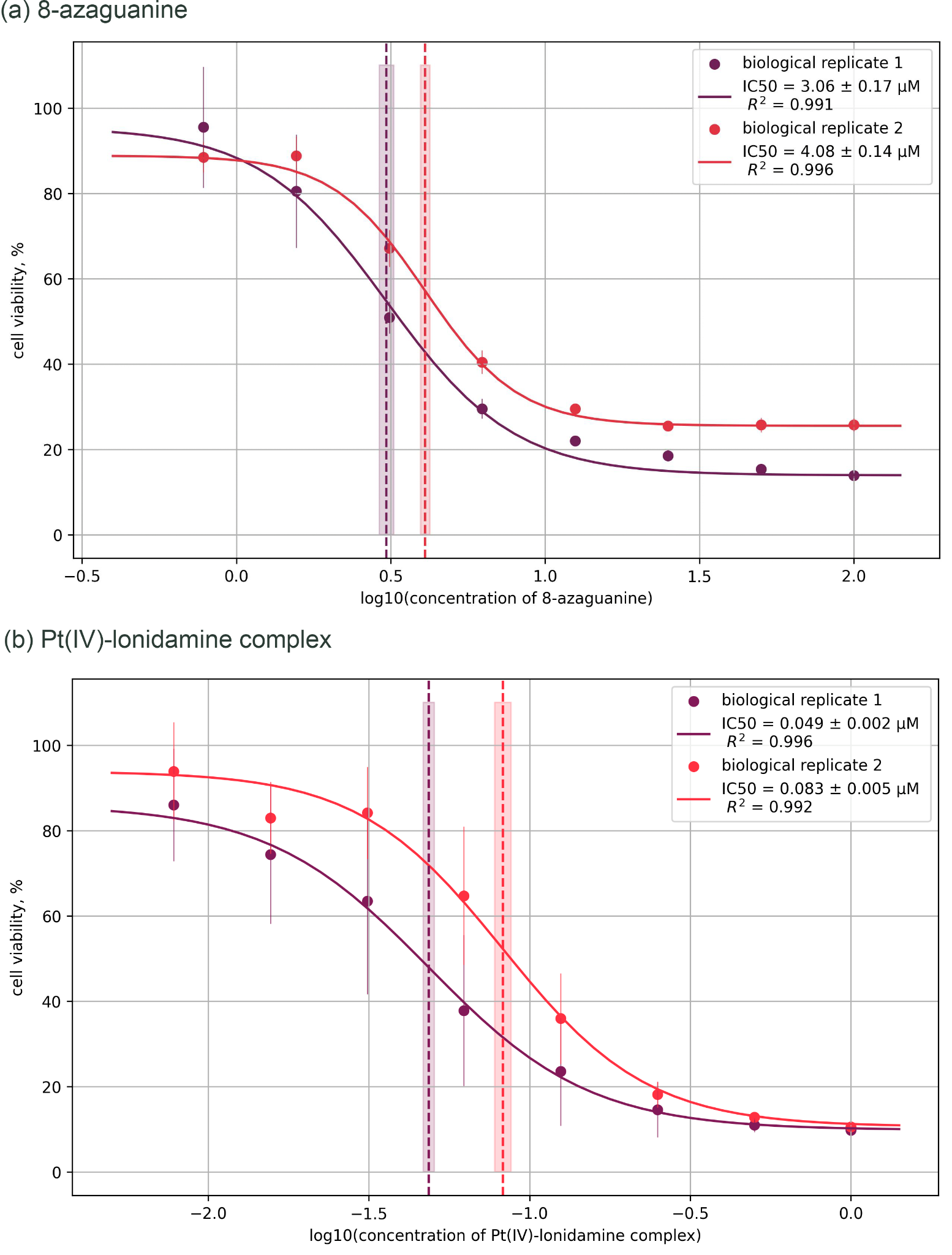

**Figure S2.** Determination of the cytotoxic effect of the (a) 8-azaguanine and (b) Pt(IV)-lonidamine complex on A2780 cell line using MTT assay. The experimental data were approximated using a dose-response curve that defines the equation for the IC50 as follow $Y = Bottom + \frac{Top-Bottom}{1+10^{(logIC50-X)*HillSlope}}$, where *X* represents the log10 concentration of the drug, *Top* represents the maximum response observed, *Bottom* represents the minimum response, and *HillSlope* describes the steepness of the family of curves.

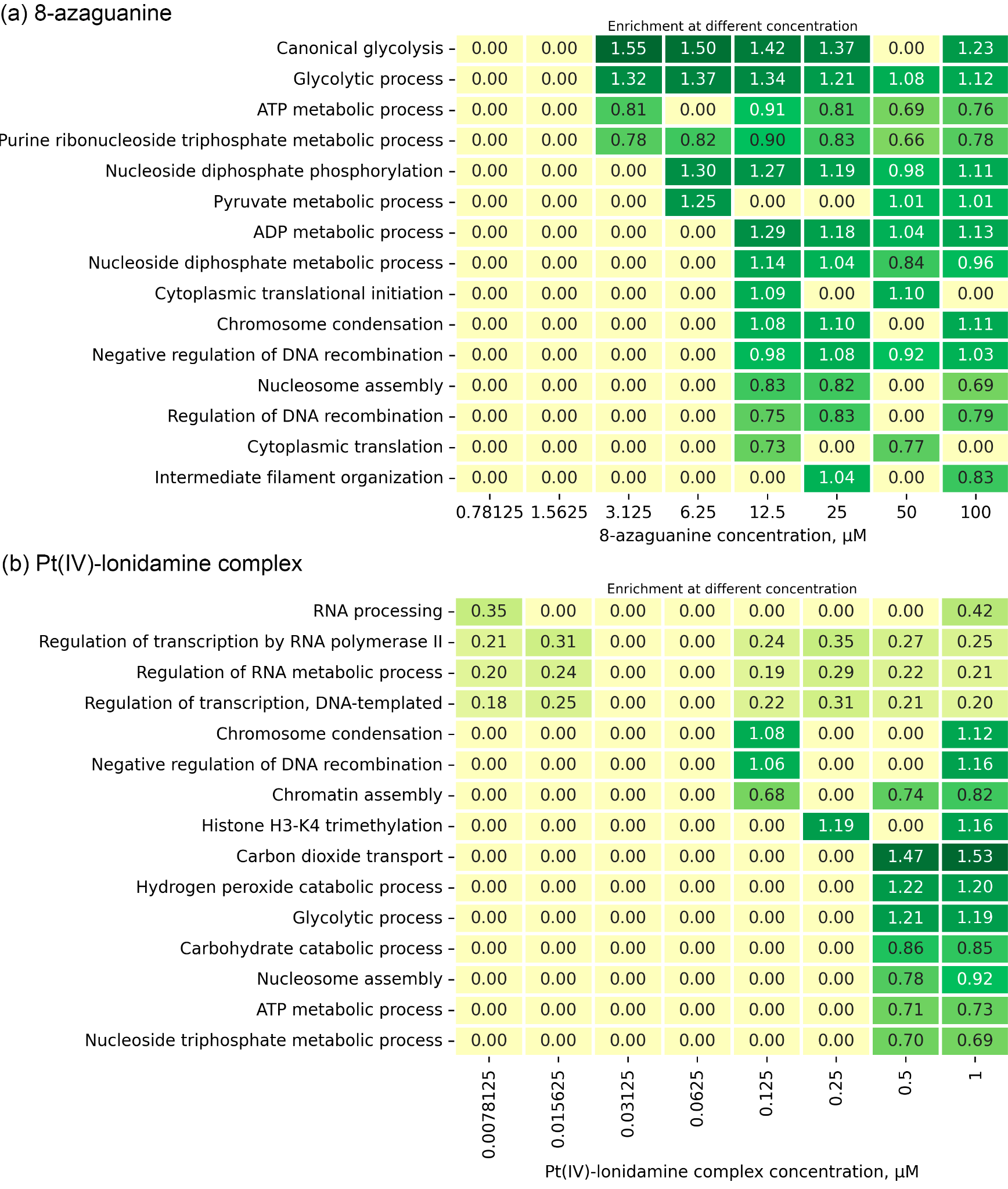

**Figure S3.** The 15 most enriched biological processes taking place in А2780 cells in response to treatment with: (a) 8-azaguanine; (b) the Pt(IV)-lonidamine complex. A criterion was established for the positive correlation between a set of biological fortifications and a set of drug concentrations.

a

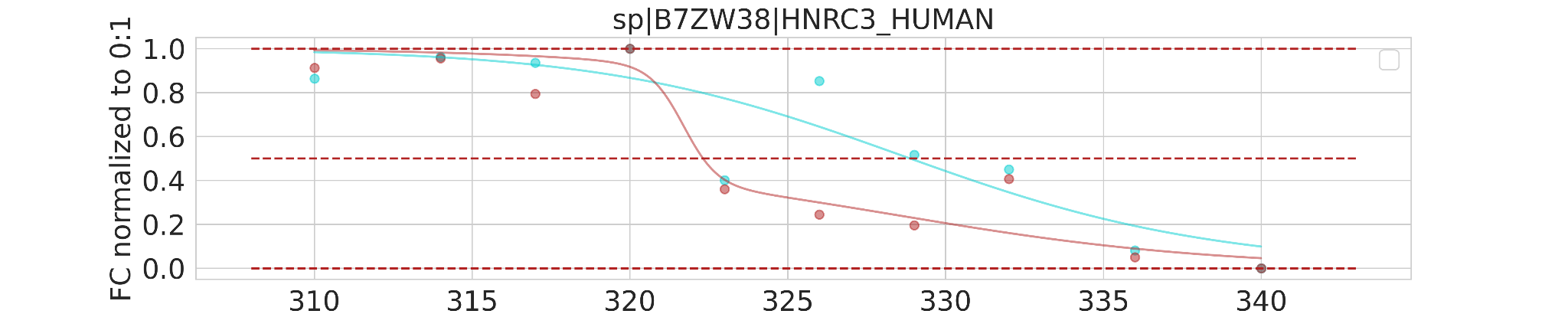

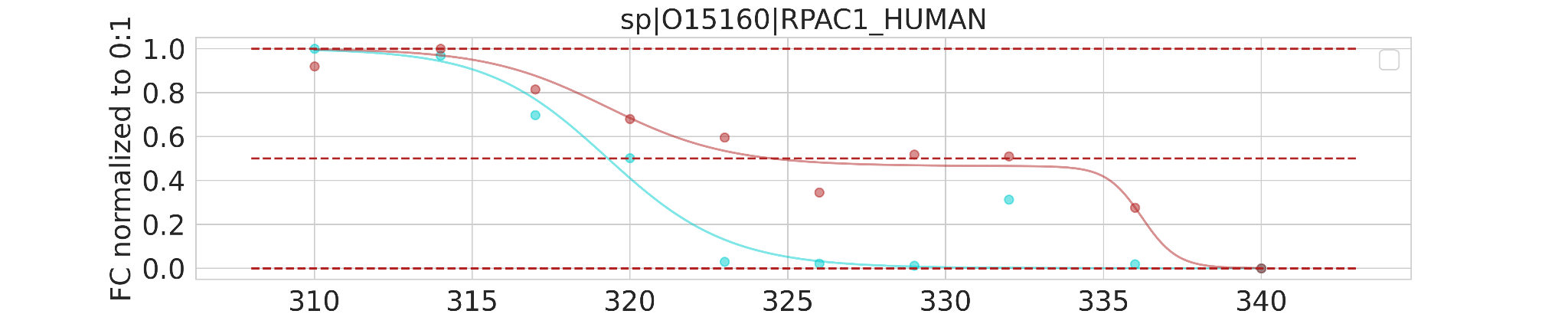

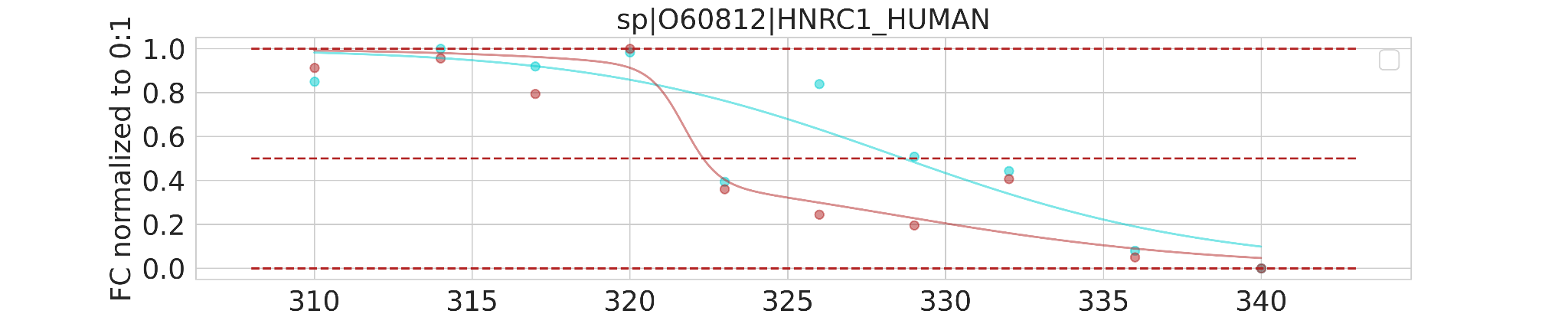

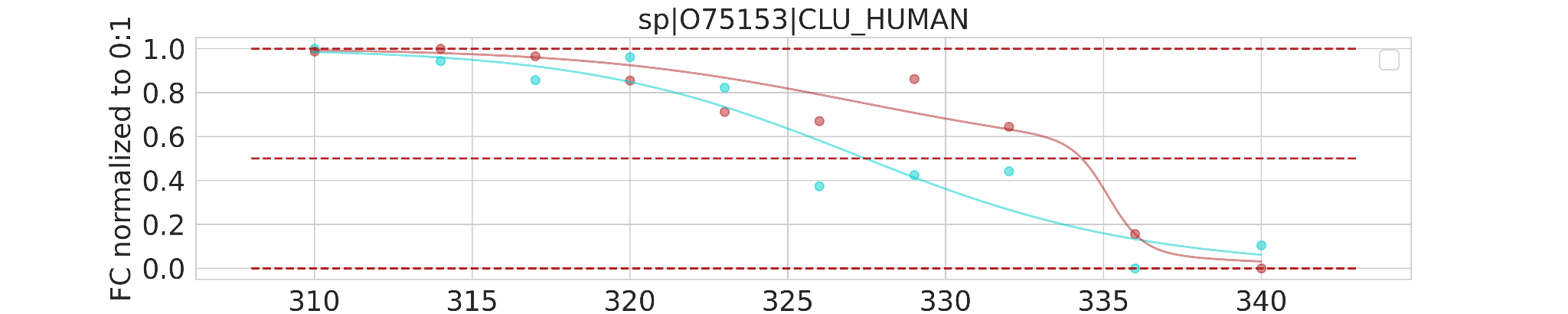

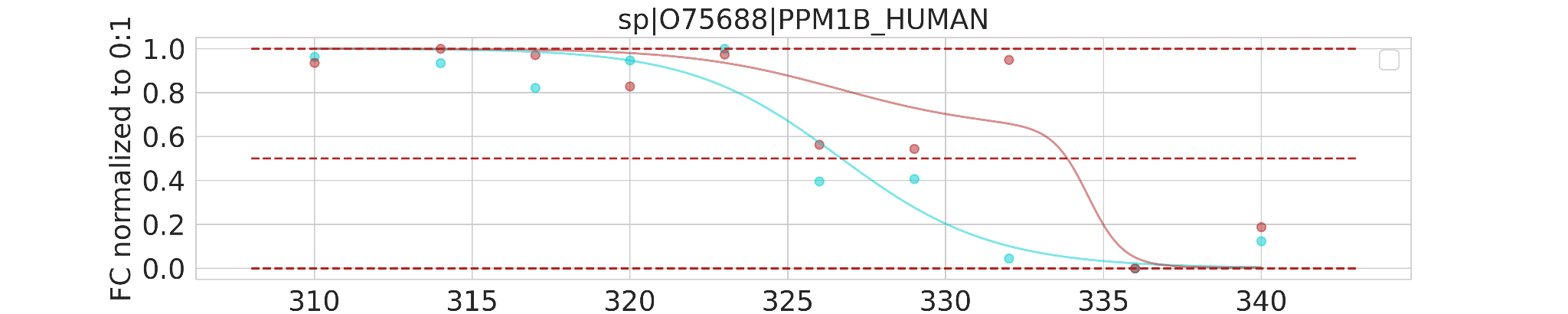

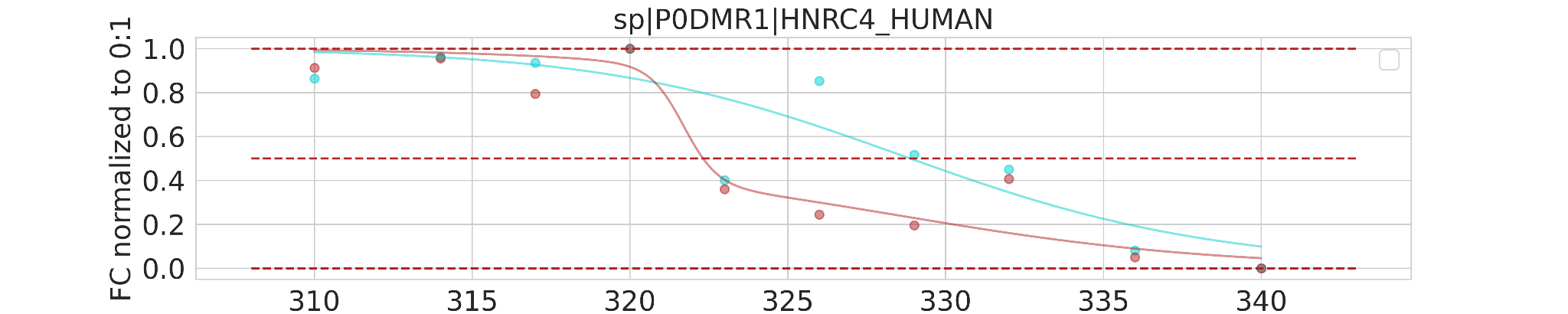

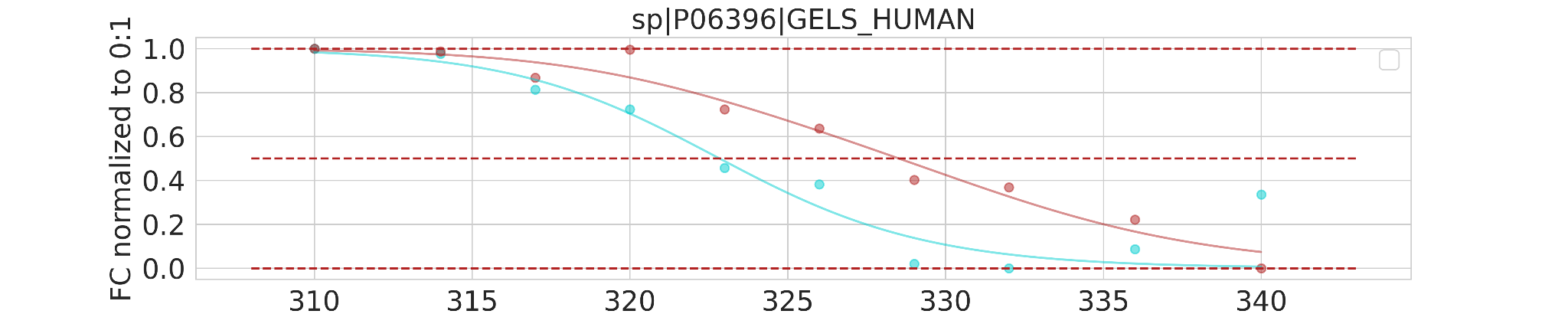

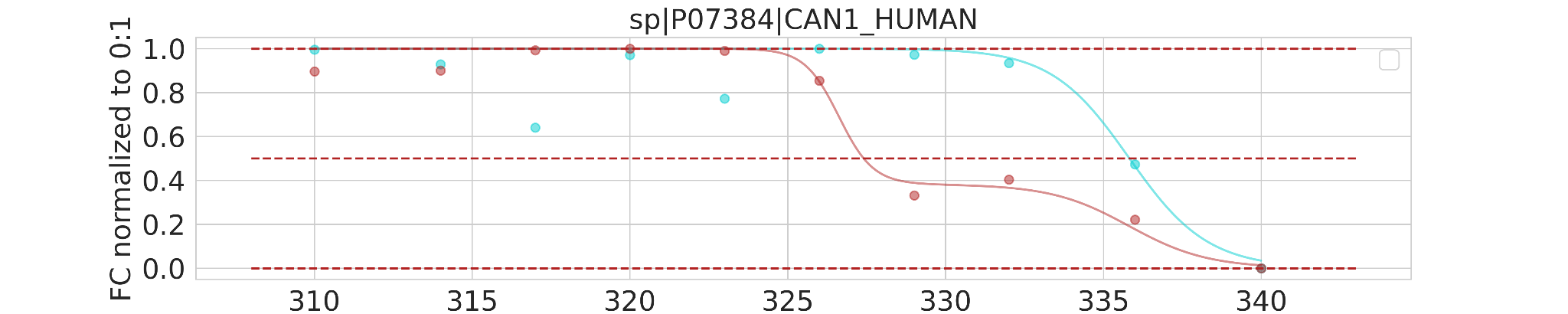

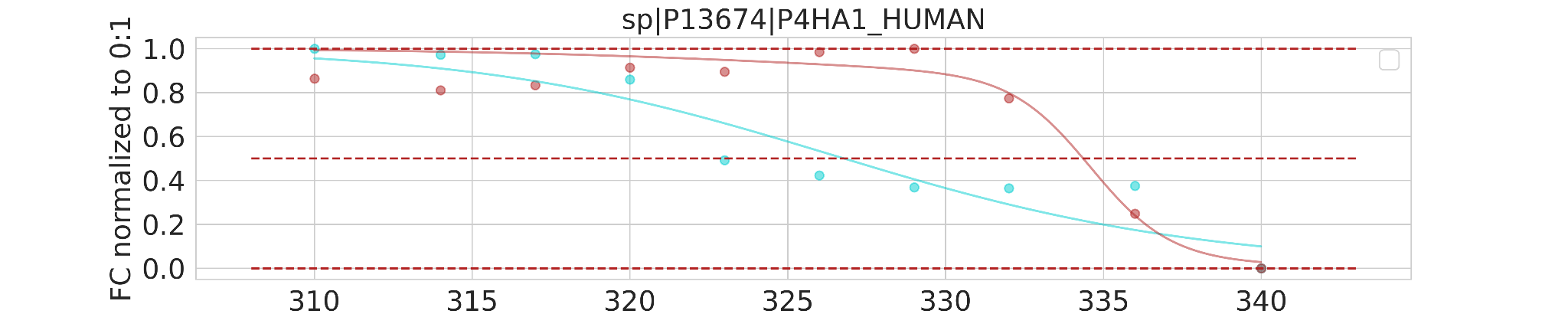

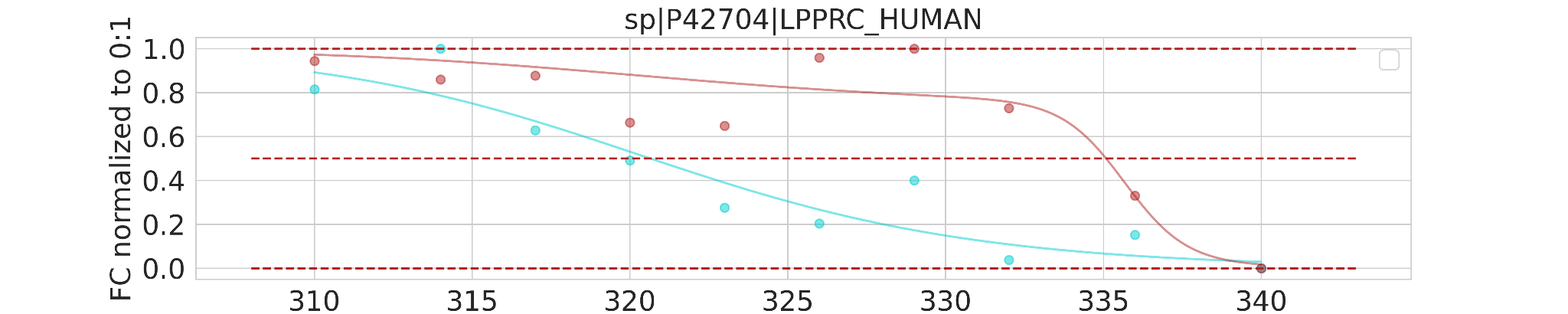

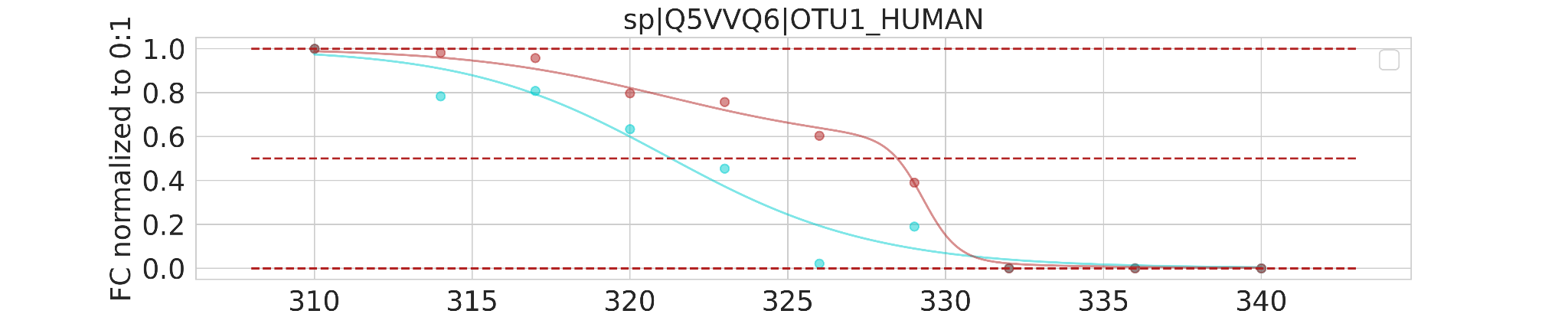

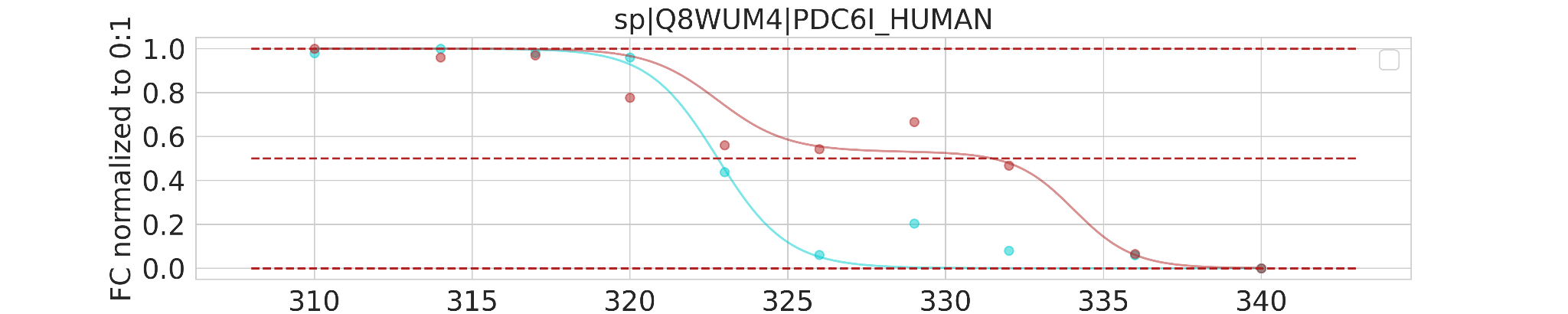

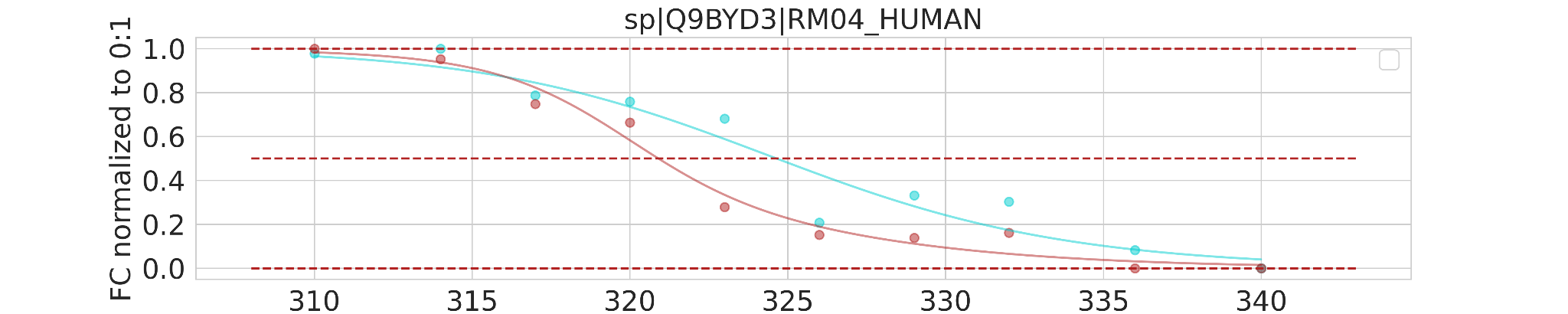

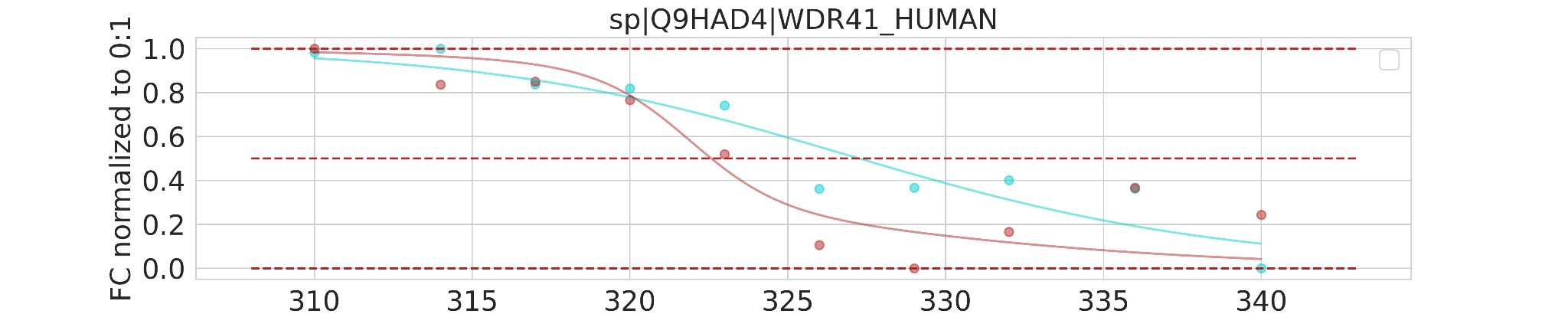

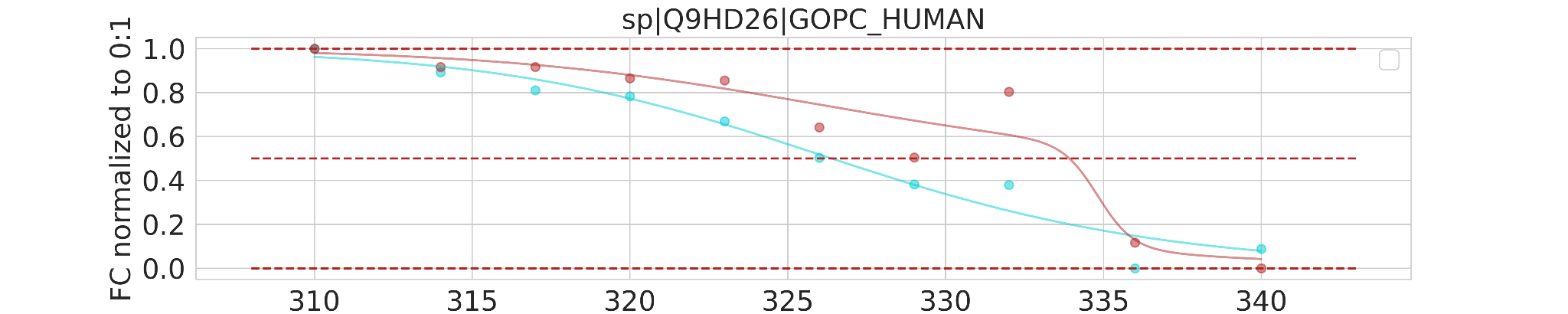

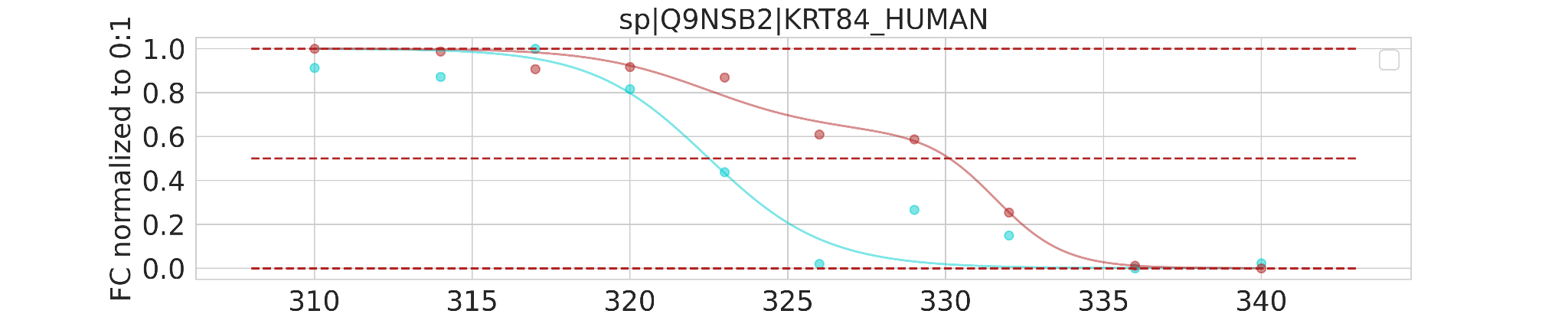

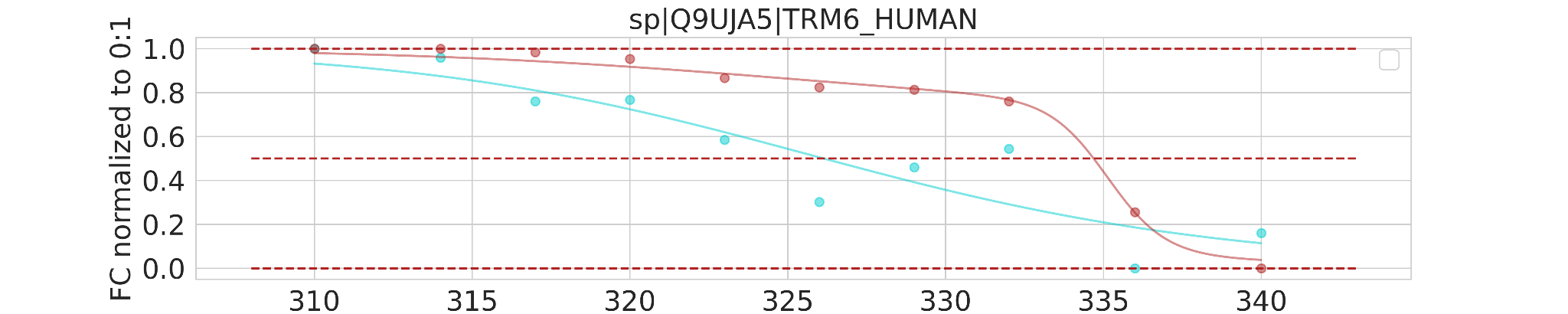

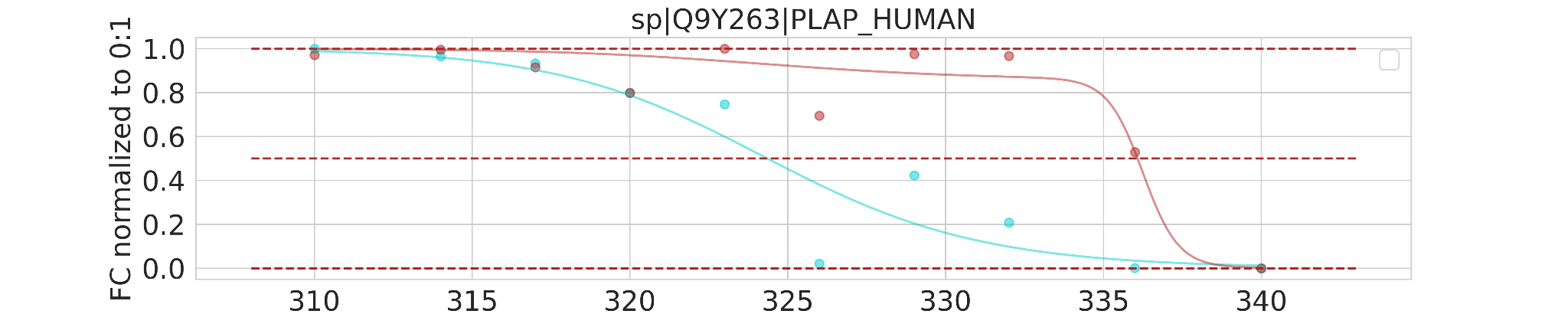

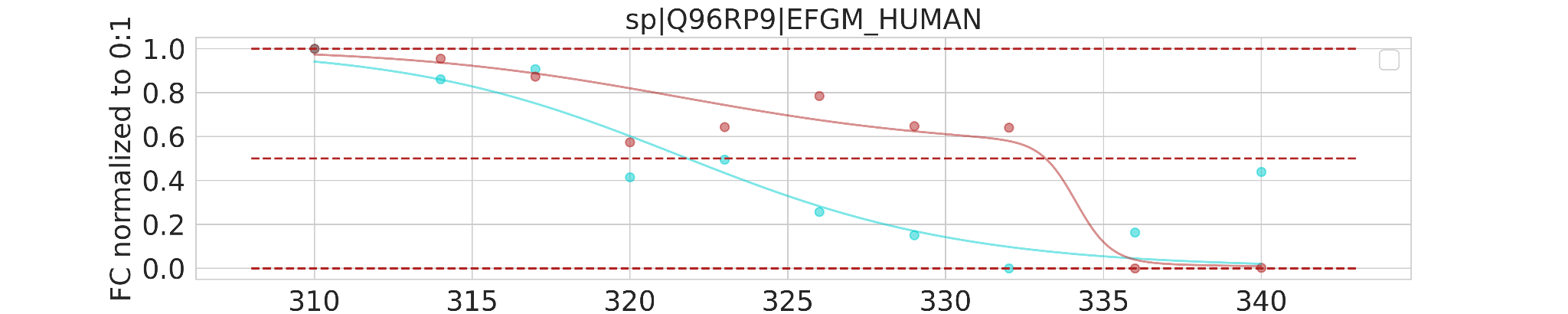

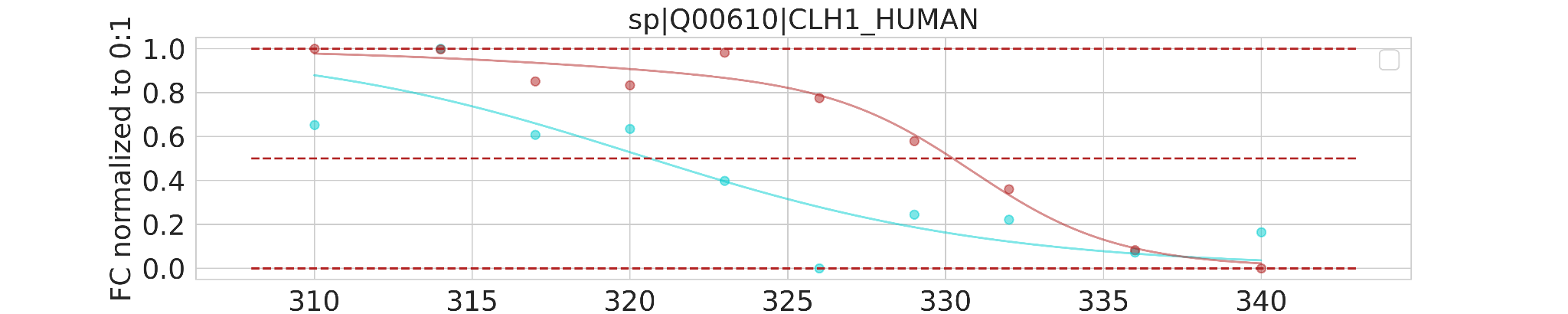

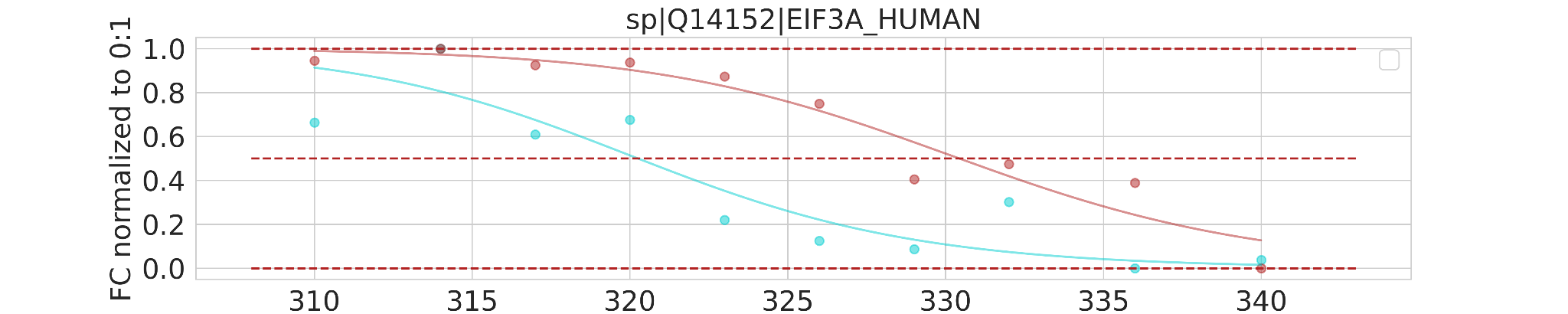

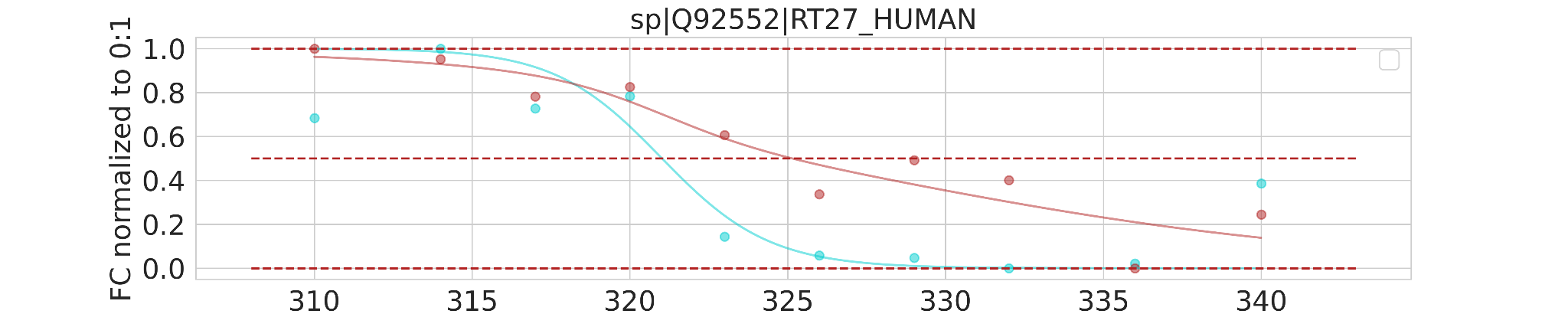

b

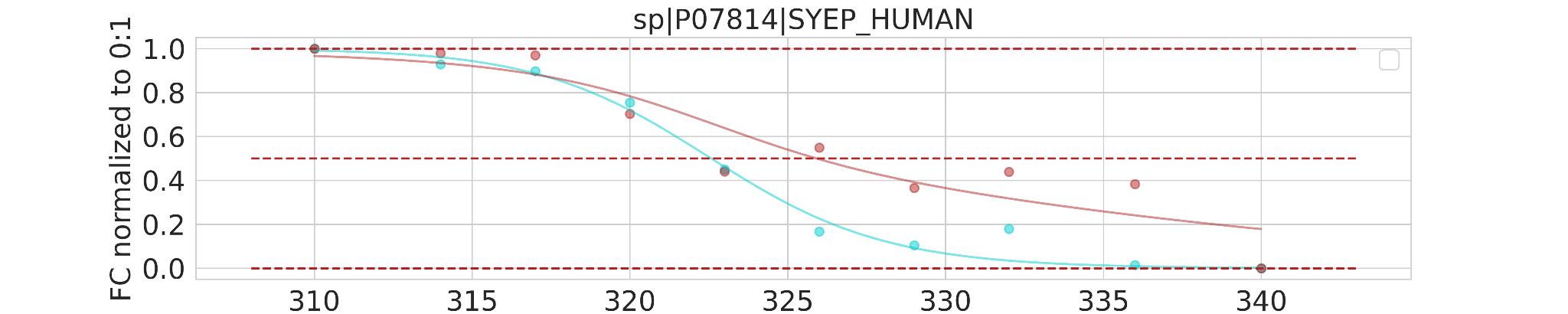

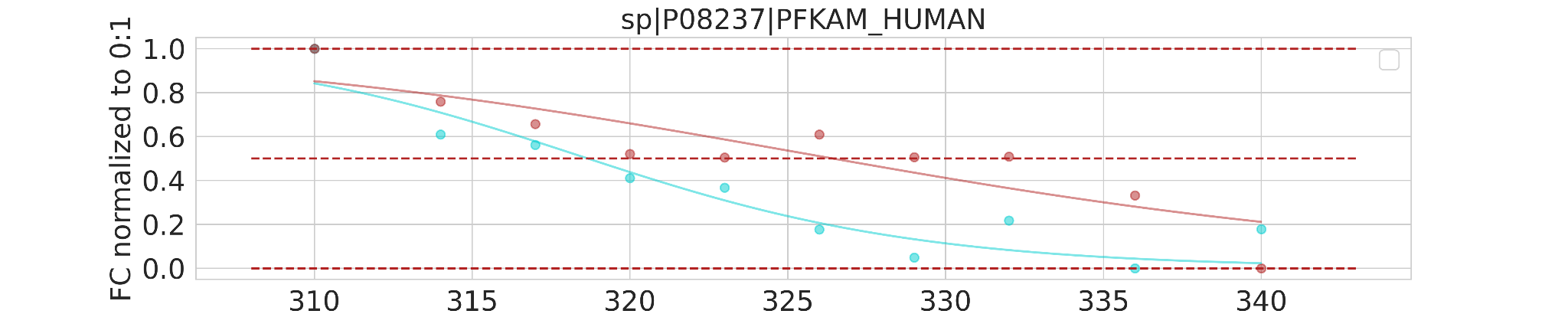

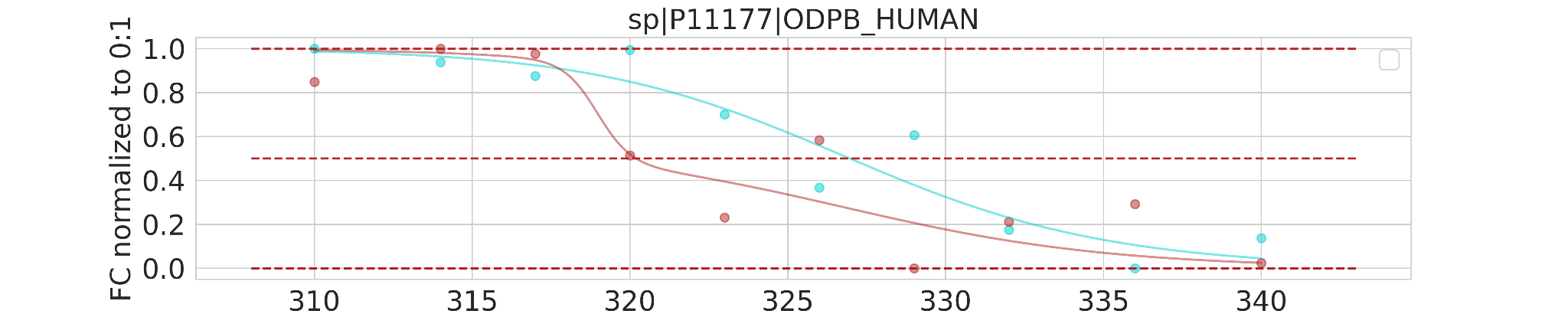

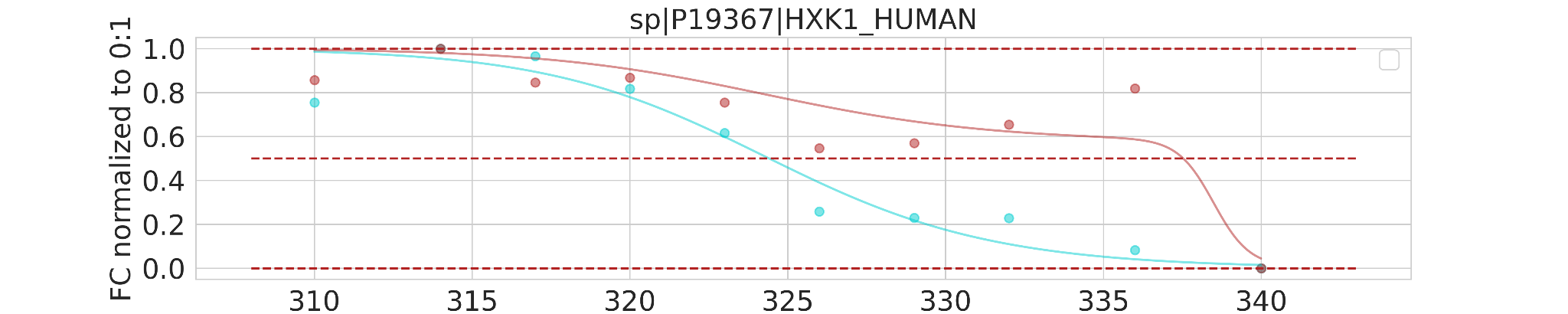

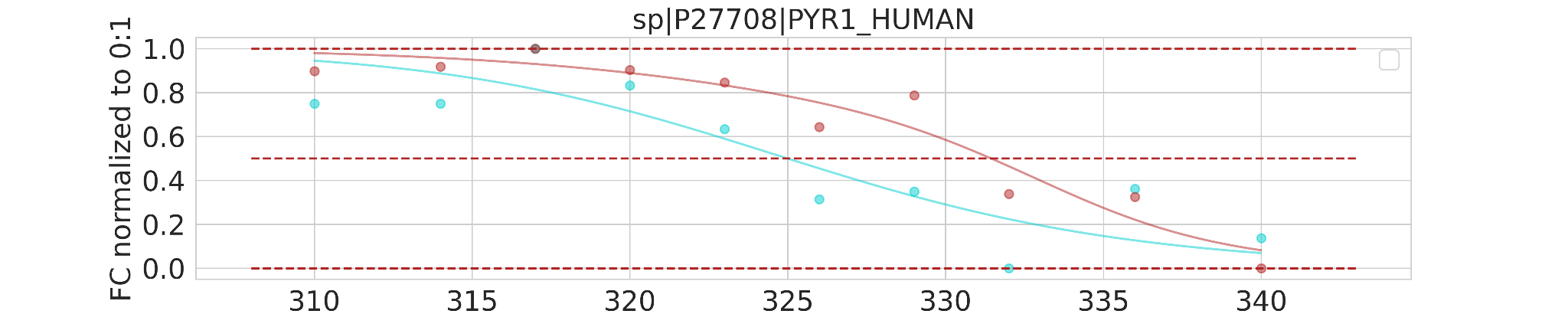

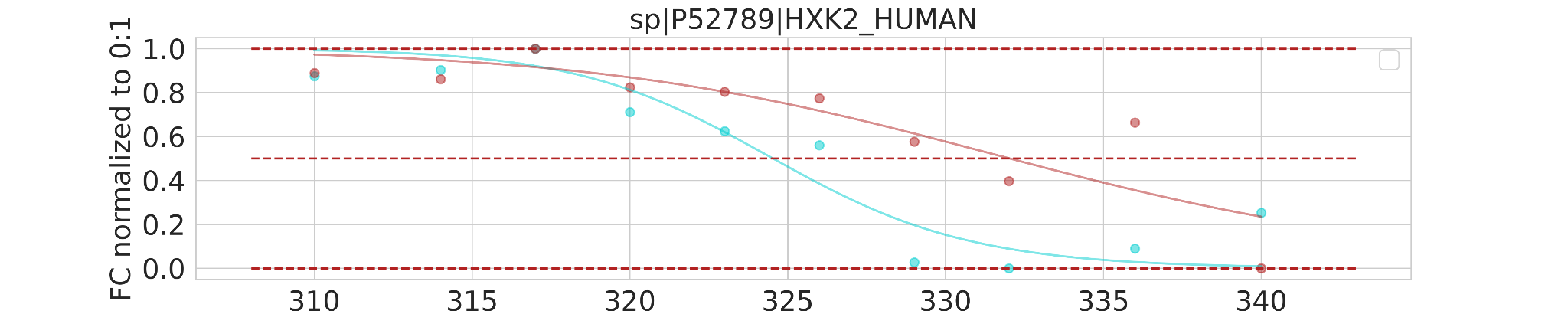

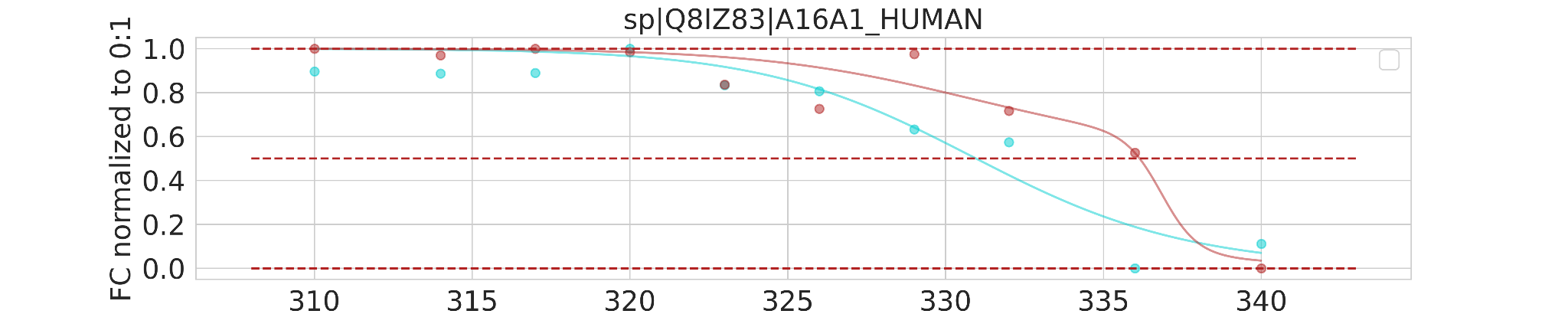

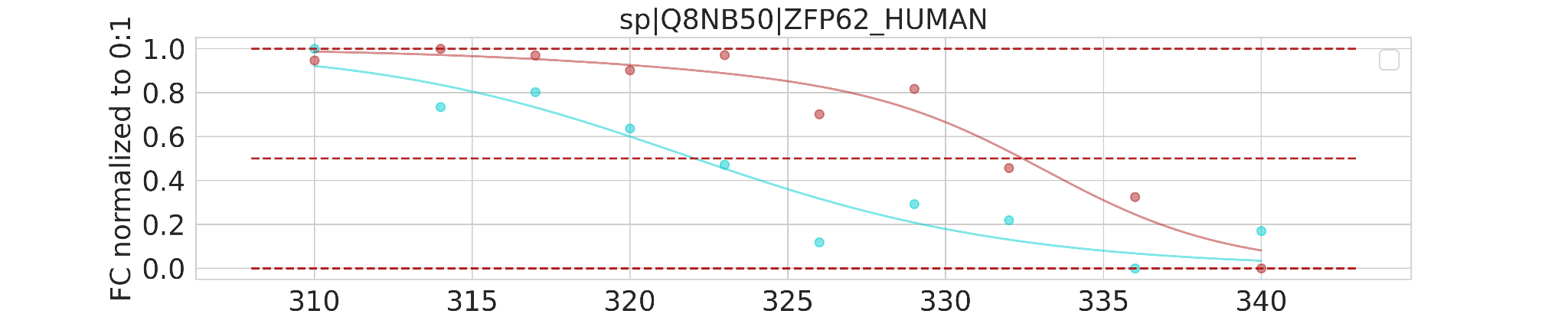

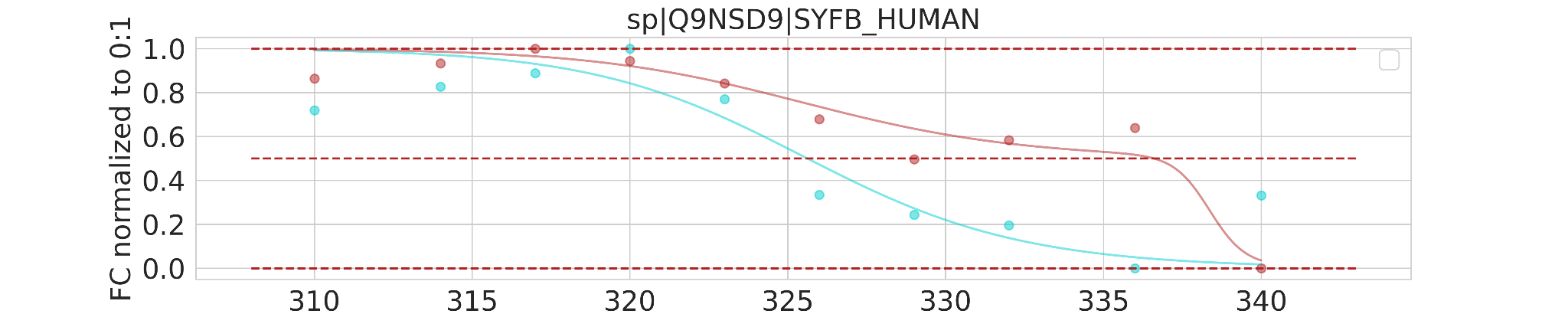

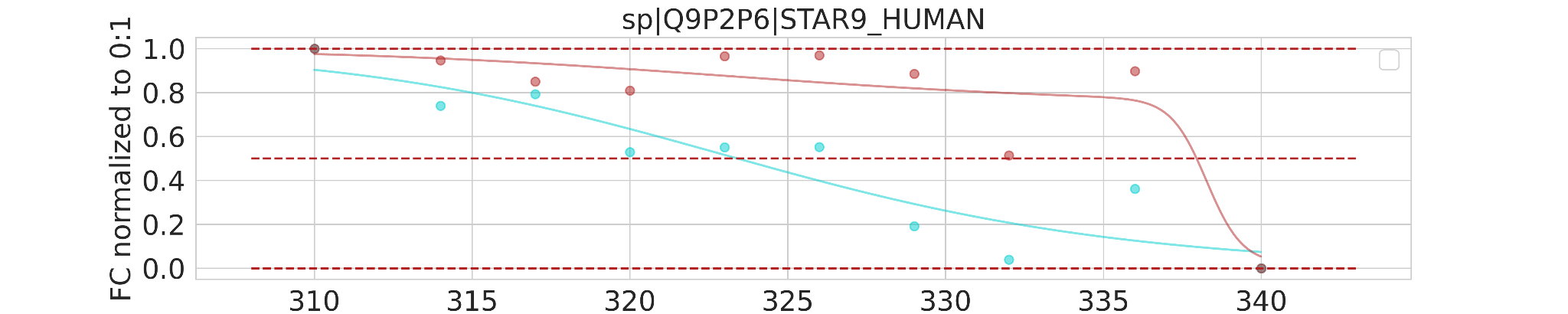

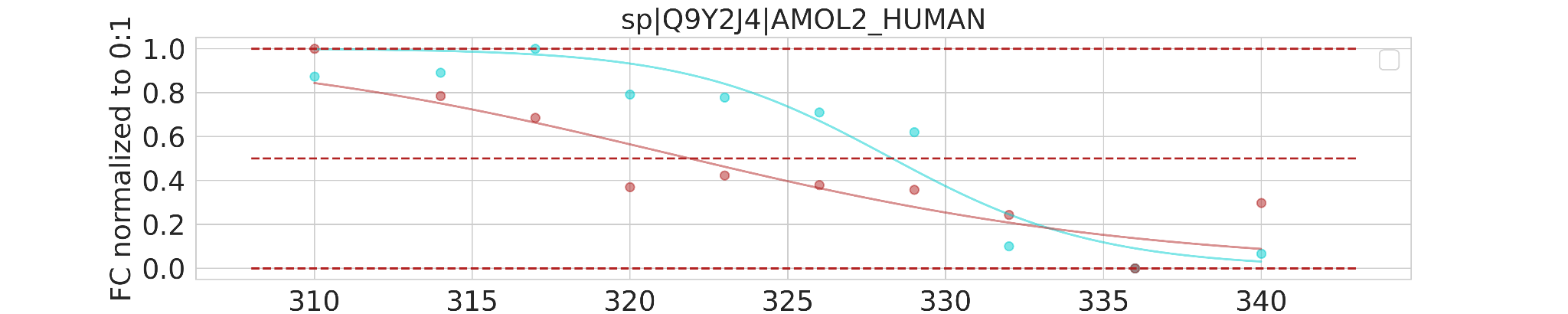

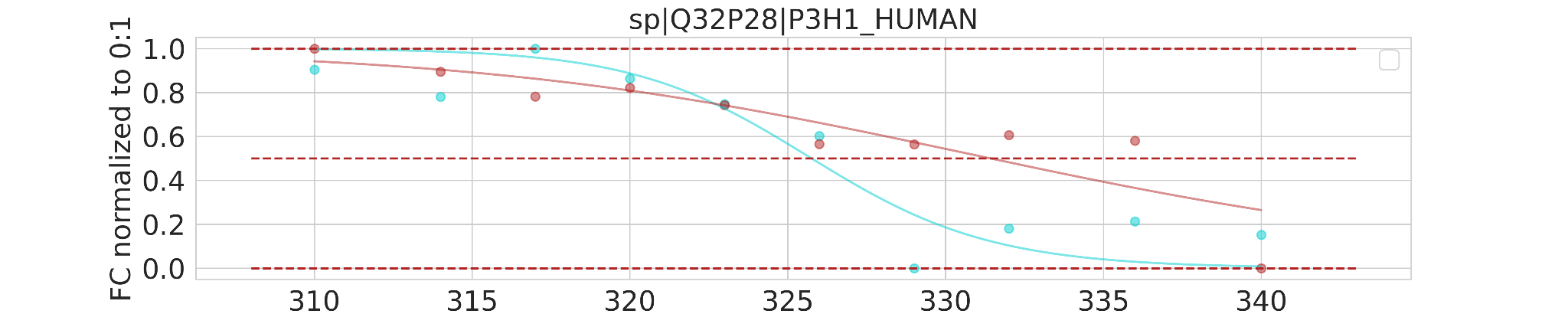

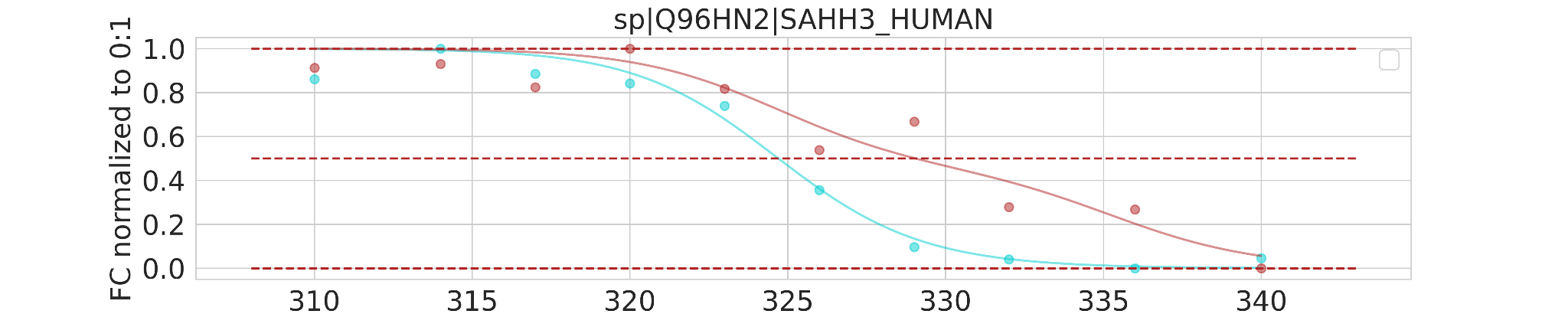

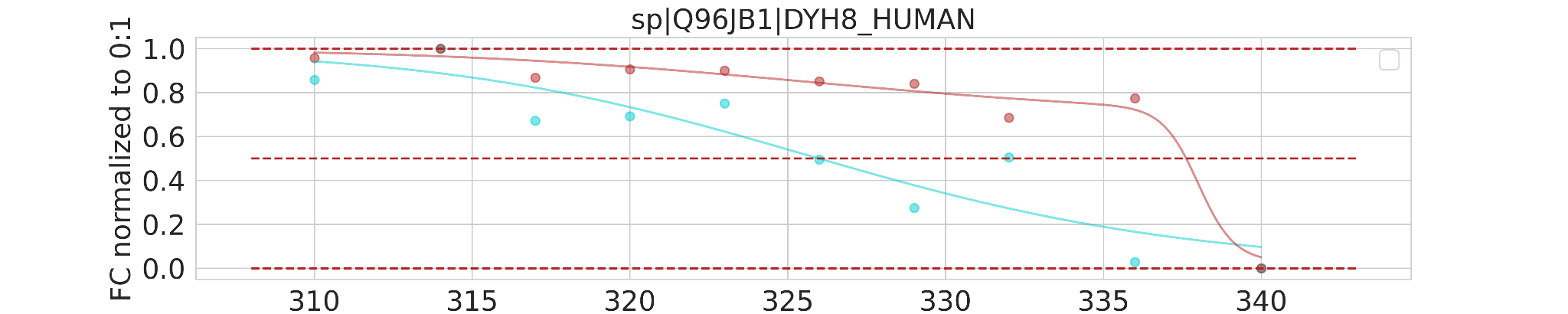

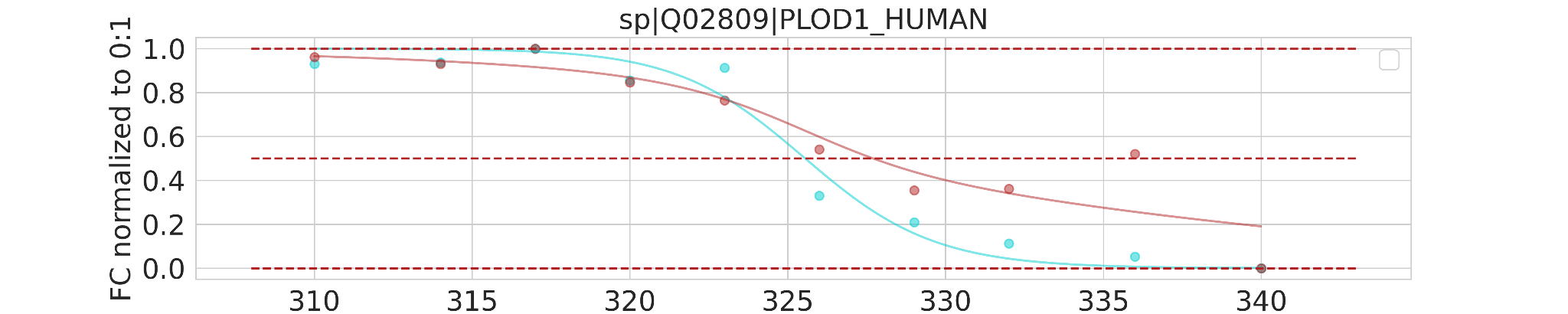

**Figure S4.** Solubility curves obtained for all selected target proteins with statistically significant deviation in the melting temperatures upon 8-Azaguanine (a) and Pt(IV)-lonidamine complex (b) treatment of the A2780 cell lysate. Blue and yellow curves correspond to control and treated samples, respectively. Dots show experimental values of protein quantification. Difference on the graphics is the difference between denaturation temperature in treated samples and control samples in Celsius degrees.
